# Nuclear SnRK1 activity delays clubroot development in *Arabidopsis* by reducing sink strength

**DOI:** 10.64898/2025.12.31.697172

**Authors:** Harshavardhanan Vijayakumar, Niel Guillaume, Lies Vandesteene, Patrick Van Dijck, Wim Van den Ende, Barbara De Coninck, Filip Rolland

**Author notes:** Correspondence: Filip Rolland, Laboratory for Plant Metabolic Signaling, Biology Department, KU Leuven, Kasteelpark Arenberg 31, 3001 Heverlee-Leuven, Belgium.

## Abstract

Clubroot, caused by the soil-borne protist *Plasmodiophora brassicae*, is a major disease of Brassica crops, resulting in severe root malformations and yield losses. While most research has centered on immune signaling and hormone dynamics, plant-pathogen interactions also dramatically reshape primary metabolism, often modifying source activity and converting infecting tissues into strong metabolic sinks. The SnRK1 (SNF1-related kinase 1) protein kinase acts as a cellular fuel gauge in plants, integrating metabolic status and environmental and developmental cues to maintain carbon and energy homeostasis. Here, we explored SnRK1-mediated quantitative resistance against clubroot disease in the related crucifer model *Arabidopsis thaliana*. Both soil and hydroponic-based disease bioassays revealed how especially increased nuclear SnRK1α1 activity antagonizes clubroot development, suggesting a pivotal role for transcriptional regulation. qRT-PCR analysis and quantification of soluble sugar contents and invertase activity in roots indicate that SnRK1 represses sucrose transporter expression as well as cell wall invertase (CWINV) expression and activity, likely limiting clubroot development by reducing sink strength. Consistently, cellular assays indicate that the recently identified SnRK1α1-targeting *P. brassicae* effector PBZF1 interferes with SnRK1α1 nuclear translocation. Our study thus corroborates that SnRK1 is a primary effector target and shows that SnRK1-mediated reprogramming of gene expression and sink activity is an effective mechanism against clubroot disease development.

## Introduction

Biotic stress caused by viruses, bacteria, fungi, oomycetes, nematodes, herbivores, and parasitic or competing plants leads to considerable and increasing crop yield losses. However, plants developed a diverse arsenal against pests and pathogens that includes but goes well beyond the physical barriers of their rigid cell walls and protective external tissues. Tremendous progress has been made in our understanding of the molecular mechanisms involved in plant-microbe interactions and plant defense with detailed insight into the dynamically evolving processes of PAMP/MAMP (pathogen or microbe-associated molecular pattern)-triggered immunity (PTI), effector-triggered susceptibility (ETS) and effector-triggered immunity (ETI or gene-for-gene resistance), in which effectors are directly or indirectly detected by resistance (R) proteins) (Chisholm, Coaker, Day & Staskawicz 2006; Jones & Dangl 2006; Dodds & Rathjen 2010). A rich repertoire of specialized (secondary) metabolites (Piasecka, Jedrzejczak-Rey & Bednarek 2015) and specific hormone-mediated signaling pathways (Pieterse, Van der Does, Zamioudis, Leon-Reyes & Van Wees 2012) further enhances plants’ ability to rapidly and systemically respond and adapt to the changing biotic environment. This conceptual framework has also been successfully applied to engineer and breed disease resistance (Dangl, Horvath & Staskawicz 2013), however, the strong selective pressure on pathogens and the specificity of R gene-mediated resistance drive rapid microbial adaptation and defense evasion, urging the development of more sustainable quantitative (multi-factorial) resistance strategies.

While research has focused predominantly on innate immunity and associated hormone signaling pathways, less attention has been given to how biotic interactions influence primary metabolism, with pathogens typically affecting source activity and transforming infected tissues into strong metabolic sinks. Moreover, primary metabolism is not only crucial for providing the cellular energy required for an effective defense (a costly process), but it is also actively involved in coordinating the plant’s response, as exemplified by the regulation of sugar transporters, cell wall invertases, programmed cell death, and pathogenesis-related (PR) genes in response to sugar supplies (Rolland, Baena-Gonzalez & Sheen 2006; Essmann *et al*. 2008; Rojas, Senthil-Kumar, Tzin & Mysore 2014; Kang *et al*. 2023; Khanna, Ohri & Bhardwaj 2023). It is becoming increasingly clear that primary carbon metabolism serves as a point of integration of very diverse developmental, environmental, as well as stress signals, with an essential regulatory role for soluble sugars such as glucose and sucrose in controlling not only plant physiology and metabolic activity but also development and stress tolerance (Rolland *et al*. 2006; Smeekens, Ma, Hanson & Rolland 2010; Ruan 2014; Li & Sheen 2016; Bezrutczyk *et al*. 2018; Kanwar & Jha 2019; Breia *et al*. 2021; Jeandet, Formela-Luboińska, Labudda & Morkunas 2022). Conversely, carbon limitation also dramatically reprograms metabolism and gene expression, limiting growth and ensuring energy homeostasis and survival (Li & Sheen 2016; Crepin & Rolland 2019; Li, Liu & Sheen 2021)

A tight control of the energy intake/use balance is crucial for all organisms, and central to this control in eukaryotic cells are the evolutionarily conserved AMPK/SNF1/SnRK1 protein kinases. These proteins typically function as hetero-trimeric complexes with a catalytic α subunit and regulatory (scaffolding) β and (adenine nucleotide binding) γ subunits, required for stability, substrate specificity, localization and activity (Baena-González, Rolland, Thevelein & Sheen 2007; Polge & Thomas 2007; Baena-González 2010; Tomé *et al*. 2014; Emanuelle *et al*. 2015; Broeckx, Hulsmans & Rolland 2016). The animal AMP-activated kinase (AMPK), yeast SNF1 (Sucrose Non-Fermenting1), and plant SnRK1 (SNF1-related kinase 1) kinase orthologs act as cellular “fuel gauges” and generally trigger activation of catabolism and repression of energy-consuming anabolic reactions when energy supply is limited. In plants, stress conditions that limit photosynthesis, respiration, or carbon allocation induce a SnRK1-dependent genome-wide transcriptional and metabolic switch via direct phosphorylation of key metabolic enzymes and the regulation of diverse transcription factors to maintain energy homeostasis (Nukarinen *et al*. 2016; Cho, Wen, Wang & Shih 2016; Broeckx *et al*. 2016; Crepin & Rolland 2019; Peixoto & Baena-González 2022). While the regulatory subunits are required for kinase activity in opisthokont organisms, transient overexpression of the plant catalytic SnRK1α subunits (encoded by *SnRK1a1/KIN10* and *SnRK1a2/KIN11* in the model plant *Arabidopsis thaliana*) is sufficient to confer increased SnRK1 activity. *Arabidopsis* plants stably overexpressing *SnRK1a1/KIN10* show an increased starvation tolerance and delayed senescence phenotype (Baena-González *et al*. 2007). More recent work confirmed a default SnRK1 activity, consistent with significant autophosphorylation and repression by sugar-phosphates (notably trehalose-6-P) rather than activation by low-energy signals (Ramon *et al*. 2019; Blanford *et al*. 2024). SnRK1 activity is also regulated by subcellular localization. The SnRK1α subunit translocates to the nucleus in response to metabolic stress to activate target gene expression and control the plant’s stress response as well as growth and development (Ramon *et al*. 2019; Crepin & Rolland 2019).

An essential role for the SnRK1 kinases in biotic stress responses is also emerging (Hulsmans, Rodriguez, De Coninck & Rolland 2016). In tobacco, for example, SnRK1 overexpression increases resistance against geminivirus infection, presumably by limiting carbon and energy supply, restricting cell proliferation, and via direct interaction with and phosphorylation of viral protein effectors (Sunter, Sunter & Bisaro 2001; Hao, Wang, Sunter & Bisaro 2003; Shen, Dallas, Goshe & Hanley-Bowdoin 2014). SnRK1 has also been linked to innate immune responses against bacterial infection (Cernadas, Camillo & Benedetti 2008; Szczesny *et al*. 2010; Kim, Vo, An & Jeon 2015) and, more recently, in diverse fungal infections across different important crop species, involving different mechanisms (Kim *et al*. 2015; Filipe, De Vleesschauwer, Haeck, Demeestere & Höfte 2018; Han *et al*. 2020; Jiang *et al*. 2020; Huang *et al*. 2023; Luo, Yu, Xiao, Zhang & Peng 2024; Cao *et al*. 2024). Finally, rapid downregulation of SnRK1 activity in source leaves after herbivory attack was linked to increased assimilate transport to roots and subsequent plant recovery (Schwachtje *et al*. 2006). Clubroot disease is caused by the soil-borne obligate biotrophic protist *Plasmodiophora brassicae,* which primarily affects crucifer species (Brassicaceae, the mustard or cabbage family). The economic impact of clubroot is substantial, causing average yearly crop yield losses of 10-15% worldwide (Dixon 2009; Donald & Porter 2014). The life cycle of *P. brassicae* - from germination of resting spores in the soil via infection of root hairs, epidermal cells and wounds by primary zoospores leading to primary plasmodium formation, to the production of secondary zoospores and secondary plasmodia causing gall or club formation as a result of hyperplasia and hypertrophy and the physiology of infected plants have been relatively well characterized, often using *Arabidopsis* as a crucifer model system. Clubroot symptoms range from wilting and chlorosis to stunted growth, root swelling (clubs, disrupting transport processes), root rot, and eventually plant death. Infected plants are also often more susceptible to secondary infections by other pathogens due to weakened root systems and compromised defenses (Devos, Vissenberg, Verbelen & Prinsen 2005; Devos *et al*. 2006; Siemens *et al*. 2006; Asano & Kageyama 2006; Dixon 2006; Kageyama & Asano 2009; Ludwig-Müller, Prinsen, Rolfe & Scholes 2009; Cao, Manolii, Strelkov, Hwang & Howard 2009). Different studies revealed a pivotal role for metabolic and carbon dynamics in the development of clubroot symptoms (Keen & Williams 1969; Ludwig-Müller *et al*. 2009). Soluble sugars accumulate within the gall sink tissue (Evans & Scholes 1995; Brodmann *et al*. 2002; Li *et al*. 2018; Kong, Li, Zhan & Piao 2022), supported by increased cytosolic invertase activity and phloem development (Siemens *et al*. 2011; Walerowski *et al*. 2018), resulting in activation of sugar-associated gene expression and processes (Schuller, Kehr & Ludwig-Müller 2014; Irani *et al*. 2018).

Here, we report on SnRK1-mediated quantitative resistance to clubroot development in *Arabidopsis*. During our research, overexpression of SnRK1α1 was independently confirmed to confer resistance in another study also showing that the *P. brassicae* RxLR-motif and zinc finger-containing effector PBZF1 directly interacts with and inhibits SnRK1 (Chen *et al*. 2021). In addition to further exploring the effect of SnRK1α1 overexpression and knockout (KO) or silencing in both the Ler-0 and Col-0 backgrounds, we investigated the role of SnRK1α1 subcellular localization on clubroot disease progression. We found that increased nuclear localization of SnRK1α1 increases resistance, suggesting a pivotal role for SnRK1-mediated transcriptional reprogramming. Consistently, cellular assays showed that the PBZF1 effector inhibits SnRK1-regulated gene expression by blocking SnRK1α1 nuclear localization, not only revealing a novel mode of action but also confirming SnRK1 as a prime effector target and SnRK1 activation as a key mechanism for defense against clubroot disease. We also developed and optimized a hydroponic system to facilitate disease scoring and enable more detailed analysis of the different stages of *P. brassicae* infection and mechanisms underlying resistance. Quantification of soluble sugars and invertase activities in combination with gene expression analyses indicates that SnRK1 repression of root CWINV expression and activity, along with reduced sugar transporter expression, diminishes sink strength which serves as an important mechanism for limiting clubroot development.

## Results

### Increased nuclear SnRK1 activity significantly hampers clubroot development in Arabidopsis

To corroborate and further explore the role of SnRK1 in *Arabidopsis* defense against clubroot development, we performed disease assays with transgenic *Arabidopsis* lines with altered SnRK1α1 expression (and thus SnRK1 activity) and altered subcellular localization in different genetic (Ler-0 and Col-0 ecotype) backgrounds. The different lines were grown in soil and inoculated with *P. brassicae* resting spores (from the e3 isolate) 14 days after sowing. Disease progression, based on gall development and shoot biomass, was then quantified 28 days post-inoculation (dpi) using an established root disease index (DI, based on symptom severity scoring) and relative shoot weight (infected/non-infected, shoot index).

In the Ler-0 background, we evaluated wild-type plants, two independent SnRK1α1 overexpression (OE) lines (5.7 and 6.5), and two independent SnRK1α1 RNA interference (RNAi) lines (1.7 and 7.8) (Baena-González *et al*. 2007). The infected wild-type Ler-0 plants had a DI of 77.38, with 19.04% small galls (classes 1 and 2) and 80.96% medium to severe galls (classes 3 and 4) (Figure 1A, B, C). The SnRK1α1 OE lines (5.7 and 6.5) displayed significantly less severe clubroot symptoms, with DIs of 53.57 and 65.48, respectively. Conversely, the SnRK1α1 RNAi lines (1.7 and 7.8) exhibited more severe clubroot symptoms, with DIs of 91.67 and 95.24, respectively (Figure 1A, B, C). Whereas in the SnRK1α1 OE lines, most infected roots were class 1 and 2 (47-68%), the SnRK1α1 RNAi lines displayed predominantly class 4 root symptoms (76-85%) (Figure 1B). Thus, *P. brassicae* infection most severely impacted the SnRK1α1 RNAi lines, also significantly affecting shoot growth, as indicated by the shoot index. In contrast, the more resistant SnRK1α1 OE lines showed significantly reduced shoot weight loss (Figure 1D), confirming previous results [65].

**Figure 1.**
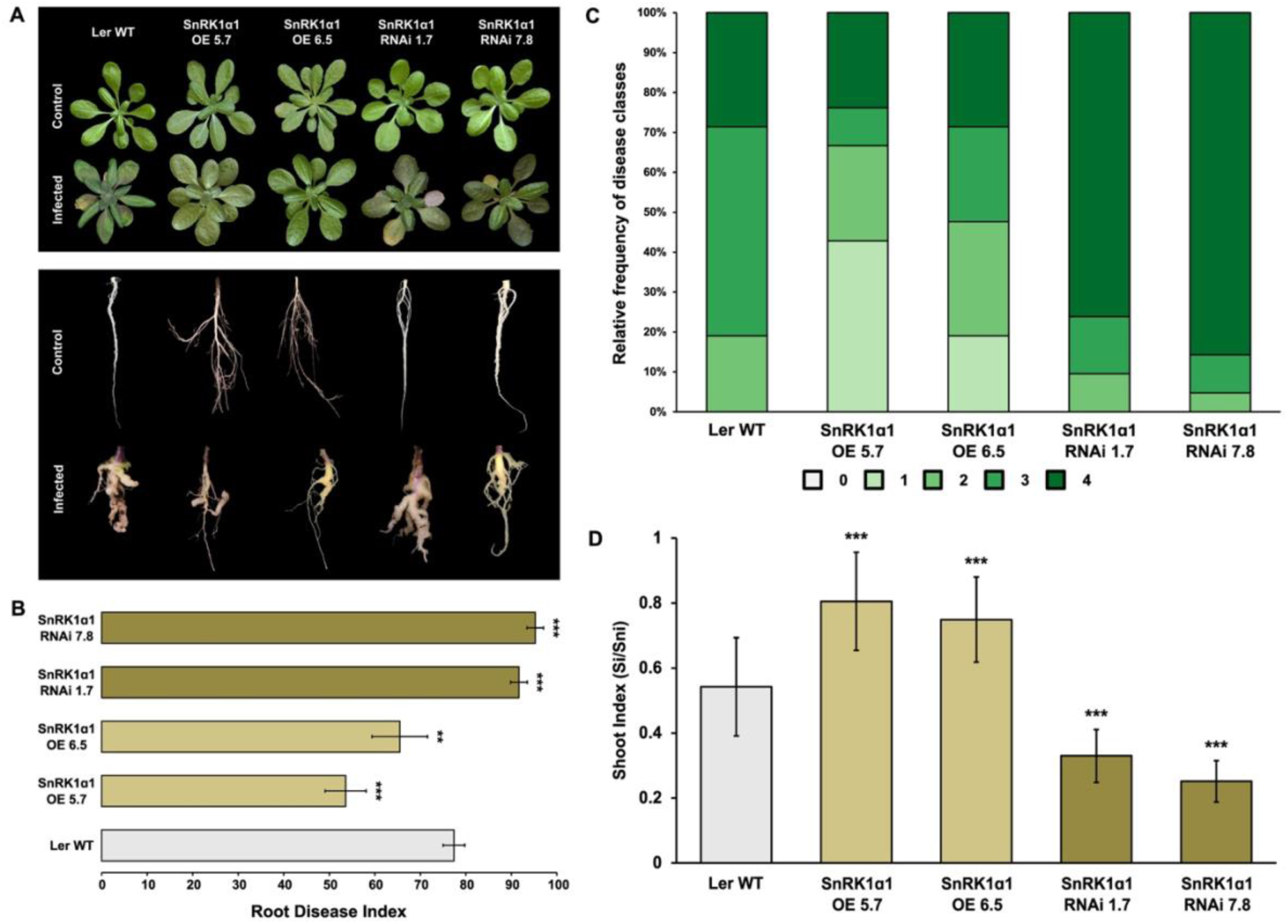
SnRK1α1 mediates resistance against clubroot disease in soil-grown Ler-0 background plants. (A) Representative shoot and root phenotypes of soil-grown infected and uninfected (control) wild-type *A. thaliana* (Ler-0 WT), two independent SnRK1α1 overexpression lines (OE 5.7 & OE 6.5), and two independent SnRK1α1 RNA interference lines (RNAi 1.7 & RNAi 7.8). (B) Root disease index (DI) quantifying overall disease severity for each genotype. (C) Relative frequency distribution of clubroot disease severity classes among the representative SnRK1α1 lines. Symptom severity was scored based on five distinct classes: Class 0, no visible symptoms; Class 1, minor swellings on minor and/or secondary roots while maintaining typical root structure; Class 2, visibly thickened primary roots, reduced fine roots and lateral roots; Class 3, significantly reduced root system with visible galls on primary and secondary roots, loss of fine roots, and occasional gall development on the hypocotyl; Class 4, roots primarily composed of a single sizeable brownish gall. (D) Relative shoot weight (shoot index, SI) was calculated for each line as the ratio of the fresh weight of infected plants to that of non-infected control plants (Si/Sni). Fourteen-day-old *A. thaliana* seedlings were inoculated with 1 mL of *P. brassicae* ‘e3’ single spore suspension (10^7^ spores/mL in 50 mM KH_2_PO_4_, pH 5.5), and disease progress was evaluated at 28 days post-inoculation (dpi). Data are the mean ± SD of three independent biological replicates, a total of 60–80 plants. Error bars represent the standard error of the means of the three replicates. Asterisks indicate a significant difference compared with the wild-type (for *** p < 0.001; two-way analysis of variance).

In the Col-0 background, wild-type plants, a single *snrk1α1* T-DNA line, and three independent SnRK1α1 OE lines (1.3, 3.1, and 7.3) were assessed for disease symptoms after clubroot infection. Wild-type Col-0 lines had a DI of 80.95, with 9.52% small galls and 90.47% medium to severe galls (Figure S1A, B, C). The *snrk1α1* line displayed a gall development similar to that of wild-type plants, with a DI of 82.41 and most infected roots in classes 3 and 4 (88.89%). Conversely, all three SnRK1α1 OE lines showed reduced gall formation, with DIs of 57.41, 60.19, and 53.70, respectively, primarily with roots in classes 1 and 2 (51-60%) (Figure S1A, B, C). Furthermore, the shoot index of the SnRK1α1 OE lines was significantly higher than those of wild-type and *snrk1α1* lines (Figure S1D).

To investigate the impact of SnRK1 subcellular localization on *P. brassicae* infection, we also evaluated *snrk1α1 snrk1α2* double KO mutants complemented with genomic fragments encoding a wildtype (unmodified) SnRK1α1 (SnRK1α1), SnRK1α1 fused to an SV40 nuclear localization signal (NLS-SnRK1α1, with increased nuclear localization), or SnRK1α1 fused to a β-subunit myristoylation motif peptide (βMYR-SnRK1α1, with cytoplasmic retention because of myristoylation and membrane association) (Ramon *et al*. 2019). While SnRK1α1-complemented plants had a DI of 72.62, with 23.81% small galls and 76.19% medium to severe galls, NLS-SnRK1α1 plants had a DI of only 51.85, with most infected roots in classes 1 and 2 (67.81%). βMYR-SnRK1α1 plants conversely showed more severe symptoms, with a DI of 90.48 and roots mainly categorized in classes 3 and 4 (Figure 2A, B, C). The shoot weight loss of βMYR-SnRK1α1 plants was consistently higher, while NLS-SnRK1α1 shoot weight was significantly less impacted compared to WT and SnRK1α1 complemented lines (Figure 2D).

**Figure 2.**
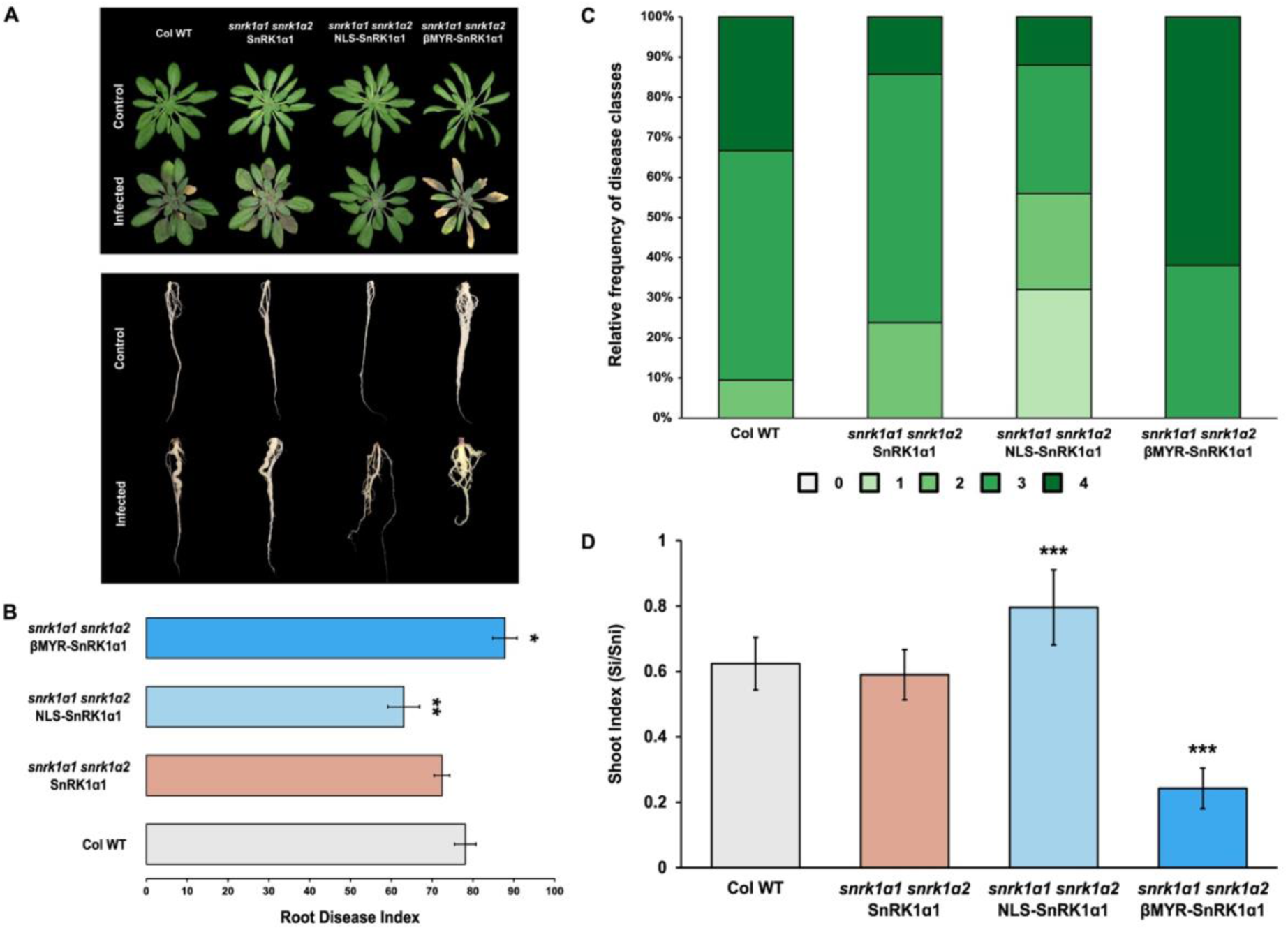
Increased nuclear SnRK1α1 localization increases resistance against clubroot disease in soil-grown plants. (A) Representative shoot and root phenotypes of soil-grown infected and uninfected (control) wild-type *A. thaliana* (Col-0 WT) and *snrk1α1 snrk1α2* double KO mutants complemented with either genomic SnRK1α1 fragments (SnRK1α1), nuclear-localized SnRK1α1 (NLS-SnRK1α1), or cytoplasmic retained SnRK1α1 (βMYR-SnRK1α1). (B) Root disease index (DI) quantifying overall disease severity for each genotype. (C) Relative frequency distribution of clubroot disease severity classes among the representative SnRK1α1 lines. Symptom severity was scored based on five distinct classes: Class 0, no visible symptoms; Class 1, minor swellings on minor and/or secondary roots while maintaining typical root structure; Class 2, visibly thickened primary roots, reduced fine roots and lateral roots; Class 3, significantly reduced root system with visible galls on primary and secondary roots, loss of fine roots, and occasional gall development on the hypocotyl; Class 4, roots primarily composed of a single sizeable brownish gall. (D) Relative shoot weight (shoot index, SI) was calculated for each line as the ratio of the fresh weight of infected plants to that of non-infected control plants (Si/Sni). Fourteen-day-old *A. thaliana* seedlings were inoculated with 1 mL of *P. brassicae* ‘e3’ single spore suspension (10^7^ spores/mL in 50 mM KH_2_PO_4_, pH 5.5), and disease progress was evaluated at 28 days post-inoculation (dpi). Data are the mean ± SD of three independent biological replicates, a total of 60–80 plants. Error bars represent the standard error of the means of the three replicates. Asterisks indicate a significant difference compared with the wild-type (for * p < 0.01; ** p < 0.001; two-way analysis of variance).

In summary, disease bioassays in soil demonstrate that SnRK1 antagonizes clubroot disease development in *Arabidopsis*, with a particularly important role for nuclear SnRK1α1 activity.

### A robust miniature hydroponic system to study clubroot disease progression and resistance mechanisms

To address the substantial limitations inherent to soil-based disease bioassays, such as heterogeneous infection and root damage and contamination during sampling (which hinder accurate disease symptom scoring and quantitative molecular analyses), we developed a standardized, cost-effective, and scalable miniature hydroponic system. This system, constructed from commercial pipette tip boxes and perforated microcentrifugation tube lids with agar plugs, offers a robust and accessible alternative for plant root-microbe interaction studies (Figure 3A, Materials and Methods).

**Figure 3.**
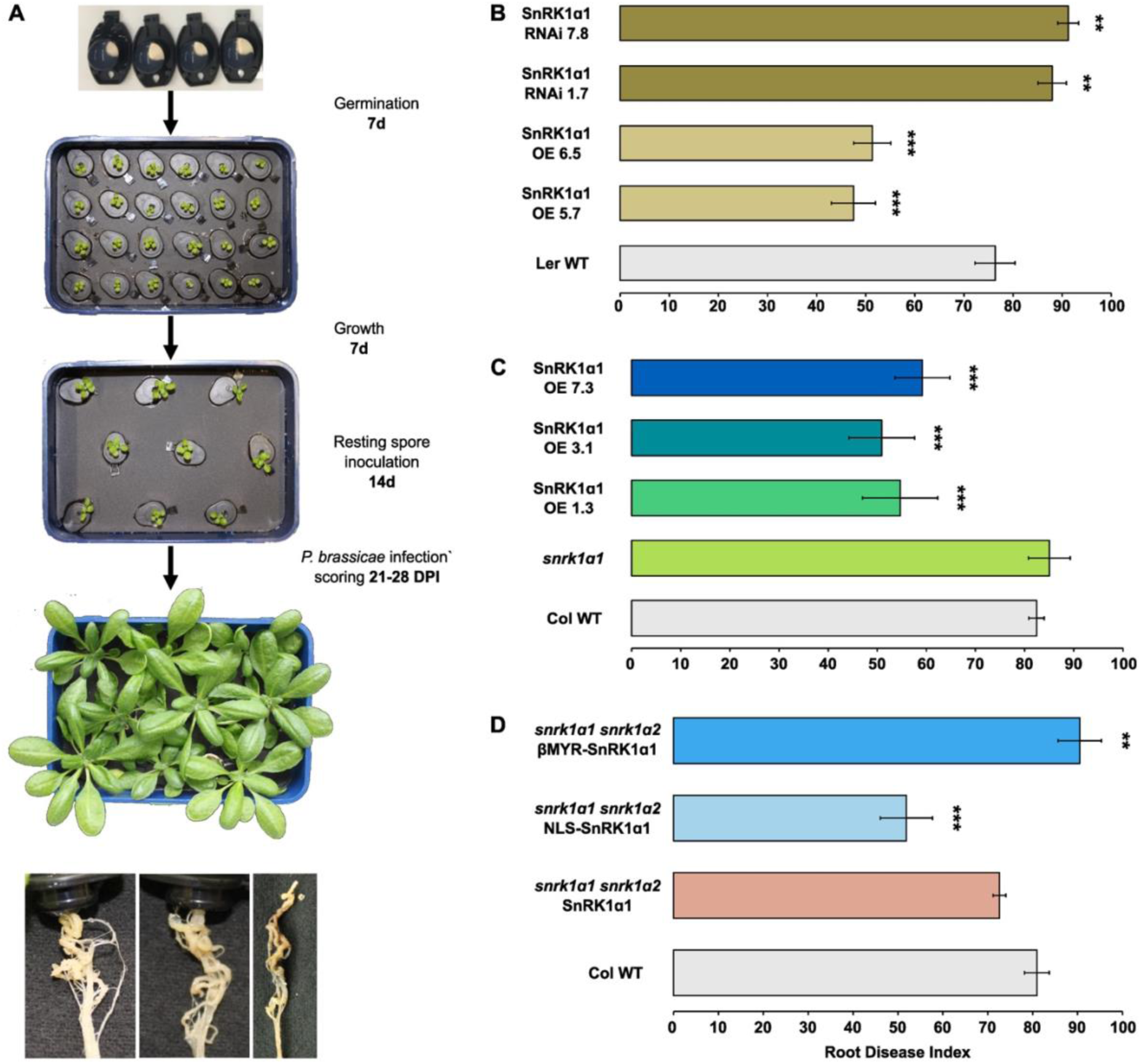
SnRK1α1 mediated resistance against clubroot disease development in a hydroponics setup. (A) A simplified hydroponic system was used for *P. brassicae* disease bioassays. Arabidopsis seeds were germinated on agar-filled microcentrifuge tube lids placed in custom 24-well holders and inserted into tip boxes containing modified ½ Hoagland medium. After 14 days of growth, seedlings were transferred to 9-well holders, and tip boxes were infected with *P. brassicae* ‘e3’ single spore isolate (10^9^ spores/mL in 50 mM KH_2_PO_4_, pH 5.5). Disease progression was assessed 21-28 dpi. Plants were grown under controlled conditions (12h light/12h dark, 21°C, 100 µmol m^-1^ s^-1^) and the medium was replenished every 2-3 days. (B-D) Root disease index (DI) quantifying overall disease severity based on the relative frequency distribution of clubroot disease severity classes among the representative SnRK1α1 lines. Data are the mean ± SD of three independent biological replicates, a total of 60–80 plants. Error bars represent the standard error of the means of the three replicates. Asterisks indicate a significant difference compared with the wild-type (for ** p < 0.01; *** p < 0.001; two-way analysis of variance).

Confirming results from the soil-based assays, Ler-0 ecotype wild-type plants produced prominent disease symptoms at 28 dpi, with 83.35% medium to severe galls (classes 3 and 4), resulting in a DI of 76.39 (Figure 3B, Suppl Figure S2A, B). SnRK1α1 OE lines exhibited a significantly lower DI (48.61 and 55.56), with a higher proportion of roots in classes 1 and 2 (55-72%). Conversely, SnRK1α1 RNAi lines had higher DIs (84.72 and 88.39) with more medium to severe galls (88.42% and 94.13%) (Figure 3B, Suppl Figure S2A, B). These effects were also reflected by the shoot index, with significantly lower and higher shoot weight loss upon infection in SnRK1α1 OE and SnRK1α1 RNAi lines, respectively (Suppl Figure S2C).

In the Col-0 background, wild-type plants had a DI of 82.41, with the majority of roots in disease classes 3 and 4 (92.59%). The *snrk1α1* line scored a similar DI of 85.01, with 96.61% of infected roots displaying medium to severe galls. In contrast, SnRK1α1 OE lines (1.3, 3.1, and 7.3) had significantly lower DIs (54.63, 50.89, and 59.17), with roots predominantly in classes 1 and 2 (56-68%) (Figure 3C, Suppl Figure S3A, B). The latter resistance was also confirmed by significantly increased shoot indices (Suppl Figure S3C).

Finally, in the modified SnRK1α1 localization lines, NLS*-*SnRK1α1 lines showed a DI of 51.85 (60.71% in classes 1 and 2), while SnRK1α1 and βMYR-SnRK1α1 lines showed Dis of 72.62 and 90.48, respectively, with a higher number of medium to severe galls (93.31% and 96.15%) (Figure 3D, Suppl Figure S4A, B). These effects were again backed up by significantly increased and decreased shoot indices in NLS-SnRK1α1 and βMYR-SnRK1α1 lines, respectively (Suppl Figure S4C).

In conclusion, our hydroponics-based clubroot disease bioassays corroborated the soil disease bioassays and validate the use of this more controlled experimental setup.

We thus started to employ this hydroponic system for subsequent analyses to better understand the biological processes and molecular mechanisms underlying SnRK1-mediated resistance. An objective way to trace and quantify infection at different stages is the quantification of pathogen DNA, in this case using *P. brassicae*-specific Internal Transcribed Spacer (ITS) regions within the ribosomal DNA (Sundelin *et al*. 2010). We analyzed representative *Arabidopsis* lines across all genotypes for the expression of *PbITS*. *PbITS* expression increased significantly faster in the wild-type and more susceptible SnRK1α1 RNAi, *snrk1α1*, and βMYR-SnRK1α1 lines compared to the SnRK1α1 OE and NLS-SnRK1α1 lines in both the Ler and Col-0 backgrounds (Figure 4), consistent with the disease assay results and confirm that nuclear SnRK1 activity hinders the proliferation of *P. brassicae*, potentially also limiting infection

**Figure 4.**
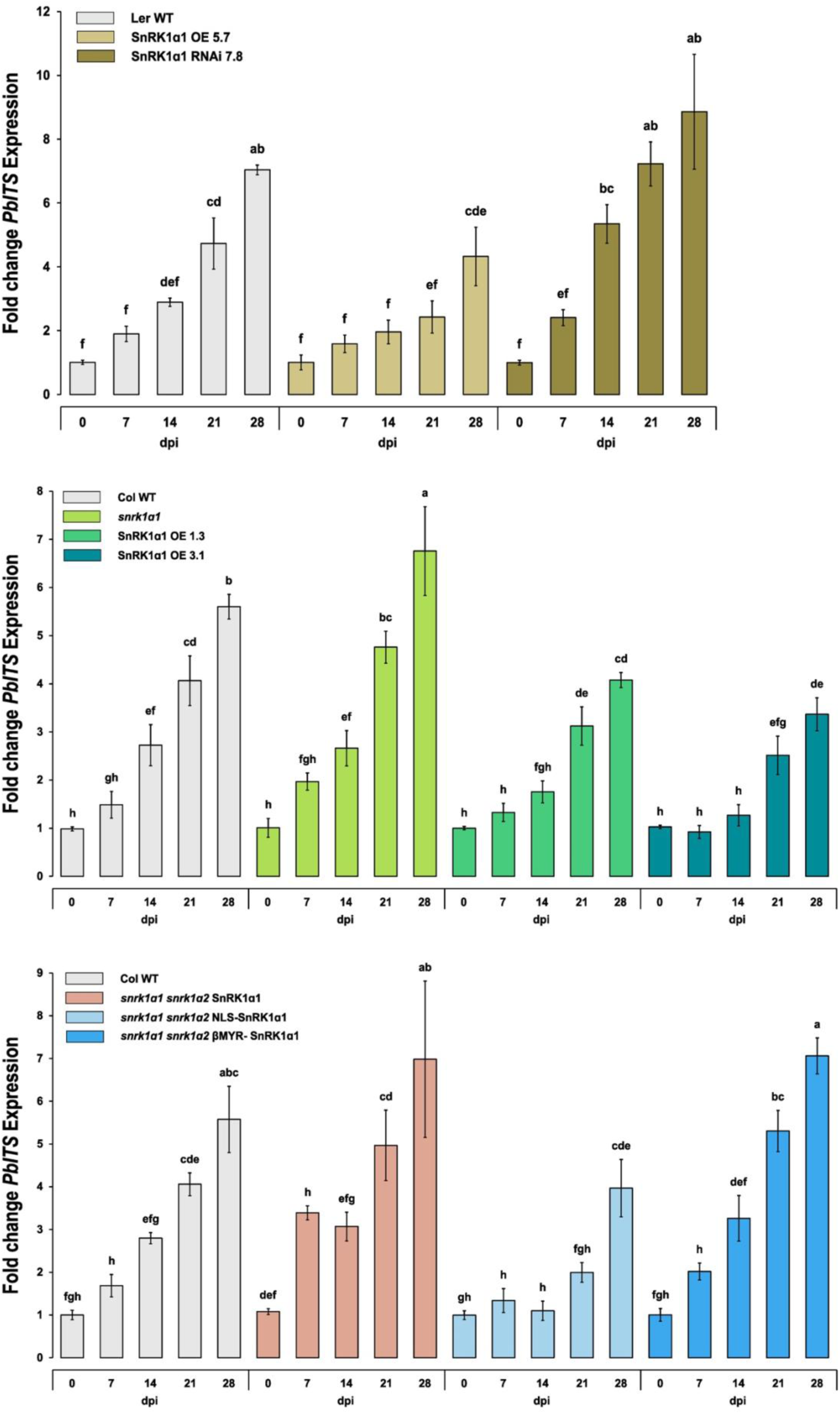
Increased nuclear SnRK1 activity reduces *P. brassicae* proliferation. (A-C) Expression of *Plasmodiophora brassicae* pathogen DNA was quantified at different time points post-inoculation (0, 7, 14, 21, and 28 dpi) among the representative SnRK1α1 lines using a *P. brassicae*-specific Internal Transcribed Spacer (ITS) region (*PbITS*). Data are the mean ± SD of three independent biological replicates, a total of 60–80 plants. Relative expression was normalized with the expression of *PbActin* and calculated against 0 dpi. Error bars represent the standard error of the means of the three replicates. Different letters represent statistically significant differences across different genotypes (p < 0.005; two-way analysis of variance).

### SnRK1 affects soluble sugar dynamics during *P. brassicae* infection

Soluble sugars are pivotal in the intricate interplay between plants and pathogens and in regulating immune responses (Liu, Song & Ruan 2022). SnRK1 maintains metabolic homeostasis, also regulating the balance of soluble sugars and adjusting their metabolism and distribution in response to carbon and energy availability, especially under stress condition. To understand the effects of sugar supply on clubroot development and SnRK1-mediated resistance, root soluble sugar (glucose, fructose, and sucrose) content was examined in the different genotypes at different time points after infection.

In the non-infected Ler-0 background, root soluble sugar content did not differ dramatically between genotypes, although the SnRK1α1 RNAi (7.8) line showed slightly increased sucrose levels. In general, soluble sugar - especially sucrose - levels increased with time and root development (Figure 5, Suppl Figure S5). In infected roots, glucose and fructose levels increased over time in wild-type and SnRK1α1 RNAi lines, peaking at 21 dpi (Figure 5A, B, Suppl Figure S5A, B). In SnRK1α1 OE roots, this increase was much more limited. Conversely, sucrose content steadily increased post-infection in all three lines, but this increase was significantly more pronounced in SnRK1α1 OE roots (Figure 5C, Suppl Figure S5C).

**Figure 5.**
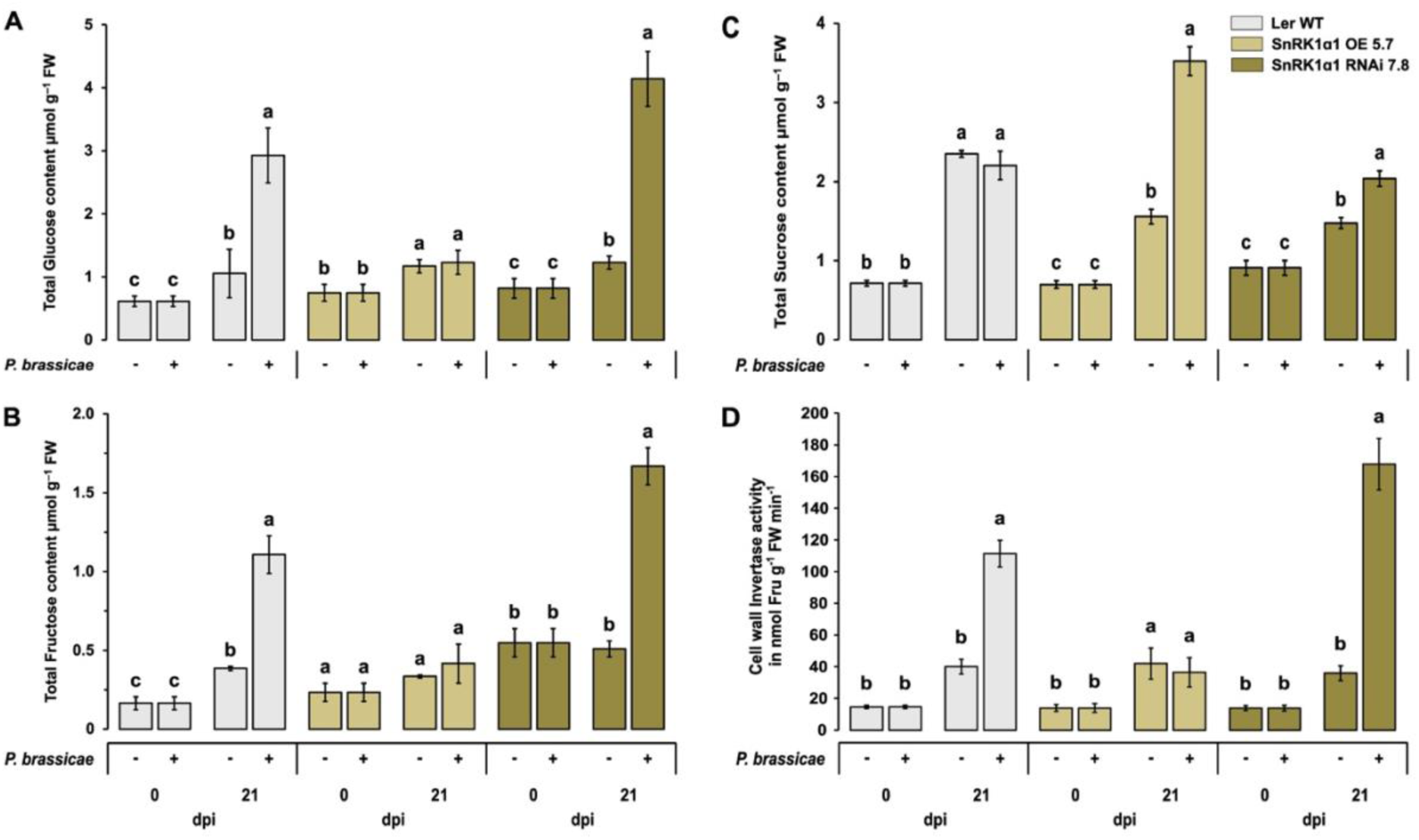
SnRK1α1 limits the impact of *P. brassicae* infection on soluble sugar dynamics and cell wall invertase activity in roots. Soluble sugar content (A) Glucose, (B) Fructose, and (C) Sucrose, and (D) Cell wall invertase activity were measured in non-infected and infected roots from hydroponically grown *A. thaliana* at 0 and 21 dpi using HPAEC-IPAD. Enzyme activity was determined by quantifying fructose production from sucrose (nmol min^–1^ g^–1^ FW). Data are the mean ± SD of three independent biological replicates, a total of 60–80 plants. Error bars represent the standard error of the means of the three replicates. Different letters represent statistically significant differences within each genotype (p < 0.005; two-way analysis of variance).

In the Col-0 background, similar effects were observed, although soluble sugar levels did not appear to increase as strongly with root development in non-infected roots. Glucose and fructose appeared to accumulate significantly more upon infection in wild-type and *snrk1α1* roots and significantly less (or not at all) in infected SnRK1α1 OE (1.3 and 3.1) roots. Conversely, sucrose accumulated significantly more in SnRK1α1 OE (1.3 and 3.1) roots (Suppl Figure S6).

When assessing the effect of SnRK1α1 localization, we observed that NLS*-*SnRK1α1 roots did not show the strong glucose and fructose accumulation observed in the other genotypes upon infection (Figure 6A, B, Suppl Figure S7A, B). Sucrose content, however, did not display the expected opposite trend as observed in the SnRK1α1 OE lines (Figure 6C, Suppl Figure S7C).

**Figure 6.**
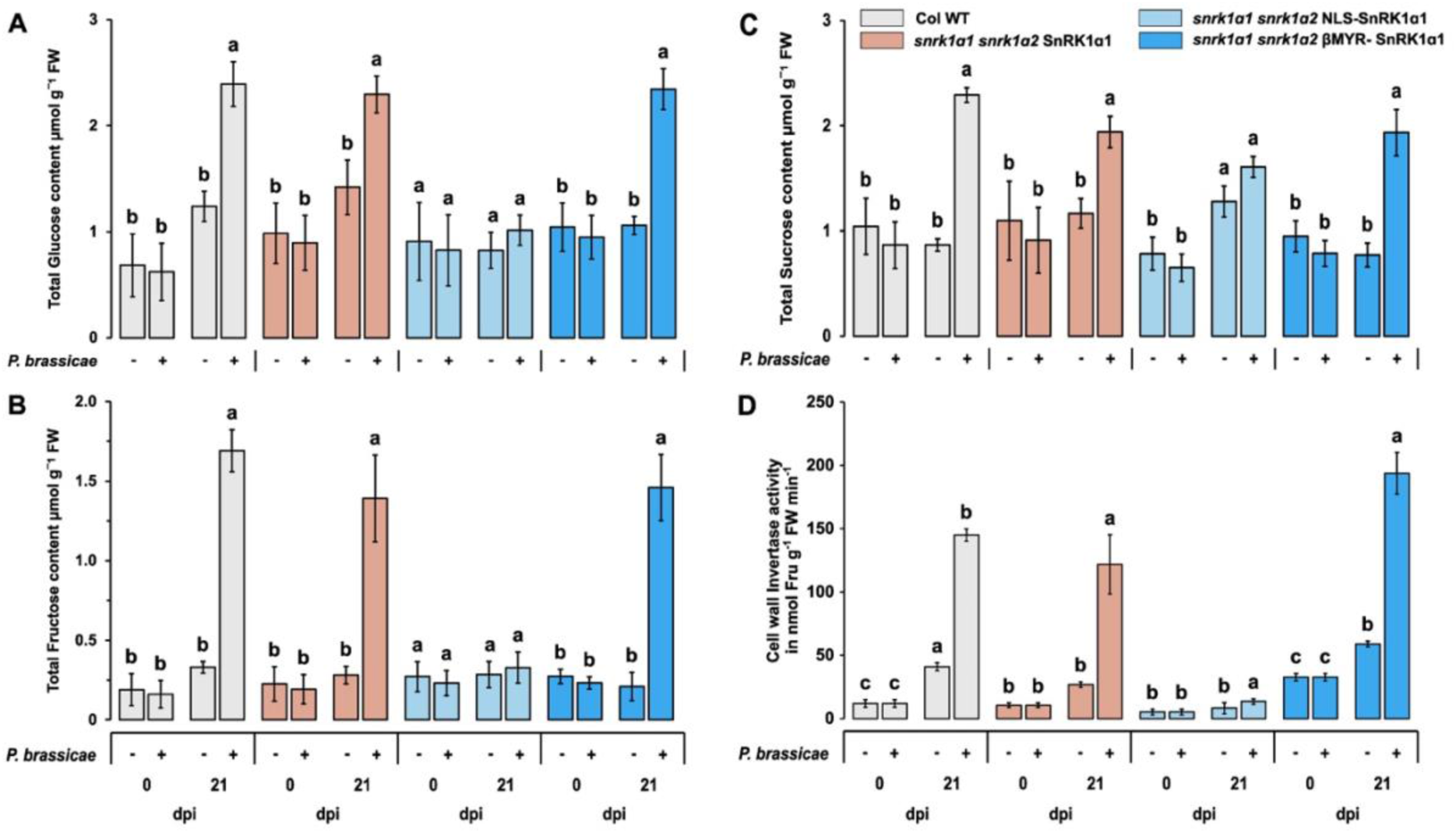
Increased nuclear SnRK1α1 localization limits the impact of *P. brassicae* infection on soluble sugar dynamics and cell wall invertase activity in roots. Soluble sugar content (A) Glucose, (B) Fructose, and (C) Sucrose, and (D) Cell wall invertase activity were measured in non-infected and infected roots of different SnRK1α1 modified-lines from hydroponically grown *A. thaliana* at 0 and 21 dpi using HPAEC-IPAD. Enzyme activity was determined by quantifying fructose production from sucrose (nmol min^–1^ g^–1^ FW). Data are the mean ± SD of three independent biological replicates, a total of 60–80 plants. Error bars represent the standard error of the means of the three replicates. Different letters represent statistically significant differences within each genotype (p < 0.005; two-way analysis of variance).

In summary, these analyses underscore the impact of *P. brassicae* infection and clubroot development on soluble sugar dynamics in *Arabidopsis* roots, with an even more pronounced increase in glucose and fructose levels in susceptible genotypes. In more resistant genotypes, this increase was limited.

### SnRK1 reduces cell wall invertase activity during clubroot infection in *Arabidopsis*

In *Arabidopsis*, sucrose is the main photo-assimilate, which is actively transported via the phloem to supply carbon and energy to developing sink organs. Enzymes such as invertases, which irreversibly hydrolyze sucrose into glucose and fructose, are thus critical in sugar partitioning (with a key role in phloem unloading, maintaining the sucrose gradient and thus sink strength) and utilization (Smeekens *et al*. 2010; Li & Sheen 2016). The soluble sugar dynamics described above suggest altered invertase activity upon clubroot infection. To elucidate the potential role of invertase activity in SnRK1-mediated resistance, we focused on the quantification of cell wall (CWINV) and vacuolar invertase (VINV) activity. CWINV activity during plant-pathogen interactions is well-documented, providing essential substrates for energy production and defense signaling (Tymowska-Lalanne & Kreis 1998; Tauzin & Giardina 2014; Veillet, Gaillard, Coutos-Thévenot & La Camera 2016). VINVs are also implicated in carbohydrate metabolism and carbon supply in response to stress conditions, but their activity’s contribution to overall sugar dynamics is typically more limited and sometimes increases susceptibility to biotic stressors (Su *et al*. 2016; Wu *et al*. 2024). The role of neutral invertase (NINV) during pathogen response is not that well documented.

In non-infected root tissues, CWINV activity was relatively low and showed a modest transient increase with time across all examined Ler-0 background lines (Figure 5D, Suppl Figure 8A). However, CWINV activity significantly yet transiently increased over time in wild-type and especially in SnRK1α1 RNAi lines, peaking again at 21 dpi (Figure 5D, Suppl Figure S8A). In SnRK1α1 OE roots, this increase was not observed. In the Col-0 background, similar effects were observed, with a strong clubroot-associated increase in CWINV activity. This increase was, however, nearly abolished in SnRK1α1 OE (1.3 and 3.1) and NLS-SnRK1α1 plants while more pronounced in βMYR-SnRK1α1 plants (Figure 6D, Suppl Figure S8B, C).

We could not detect clear differential patterns in VINV activity in the Ler-0 background lines, although all genotypes appeared to increase VINV activity upon clubroot infection (Suppl Figure S9). In the Col-0 background, this effect was not observed. In the Col-0 wild type, infection even reduced VINV activity at earlier time points. In βMYR-SnRK1α1 roots, VINV activity was consistently higher under all conditions and time points (Suppl Figure S9C).

Our analyses thus revealed clear effects on CWINV activity, consistent with the soluble sugar dynamics. While clubroot infection enhances CWINV activity, strengthening root sink capacity and carbon import into developing galls, elevated nuclear SnRK1 activity represses sucrose hydrolysis and thus carbon and energy availability for *P.brassicae*.

### SnRK1 affects *CWINV* gene expression

To elucidate how SnRK1 alters invertase activity, we examined the transcriptional regulation of two cell wall invertases, previously reported to be upregulated upon clubroot infection (Tymowska-Lalanne & Kreis 1998; Su *et al*. 2016; Veillet *et al*. 2016). The expression of *CWI1* and *CWI5* (possibly a fructan exohydrolase rather than an invertase; (Van den Ende, Lammens, Van Laere, Schroeven & Le Roy 2009)) was assessed in roots at different time points after infection using qRT-PCR (Figure 7, Suppl Figure S10, S11, S12). In the Ler-0 background, *CWI1* was significantly but transiently upregulated after infection, and this effect was significantly affected in SnRK1α1 OE (5.7) plants (showing basal levels at 21 and 28 dpi). This was even more pronounced in SnRK1α1 RNAi (7.8) plants (Figure 7A), consistent with CWINV activity. *CWI5* expression also transiently increased after infection, but no obvious differences were observed between genotypes (except for increased expression at 0 dpi in SnRK1α1 RNAi 7.8) (Figure 7B). In the Col-0 background, a strong clubroot-induced increase in both *CWI1* and *CWI5* expression was observed in both wild-type and *snrk1α1* plants (Suppl Figure S11). This induction was significantly affected in SnRK1α1 OE (1.3 and 3.1) and NLS-SnRK1α1 plants, while more pronounced in βMYR-SnRK1α1 plants (Figure 7F, 7G, Suppl Figure S11, S12), again consistent with CWINV activity.

**Figure 7.**
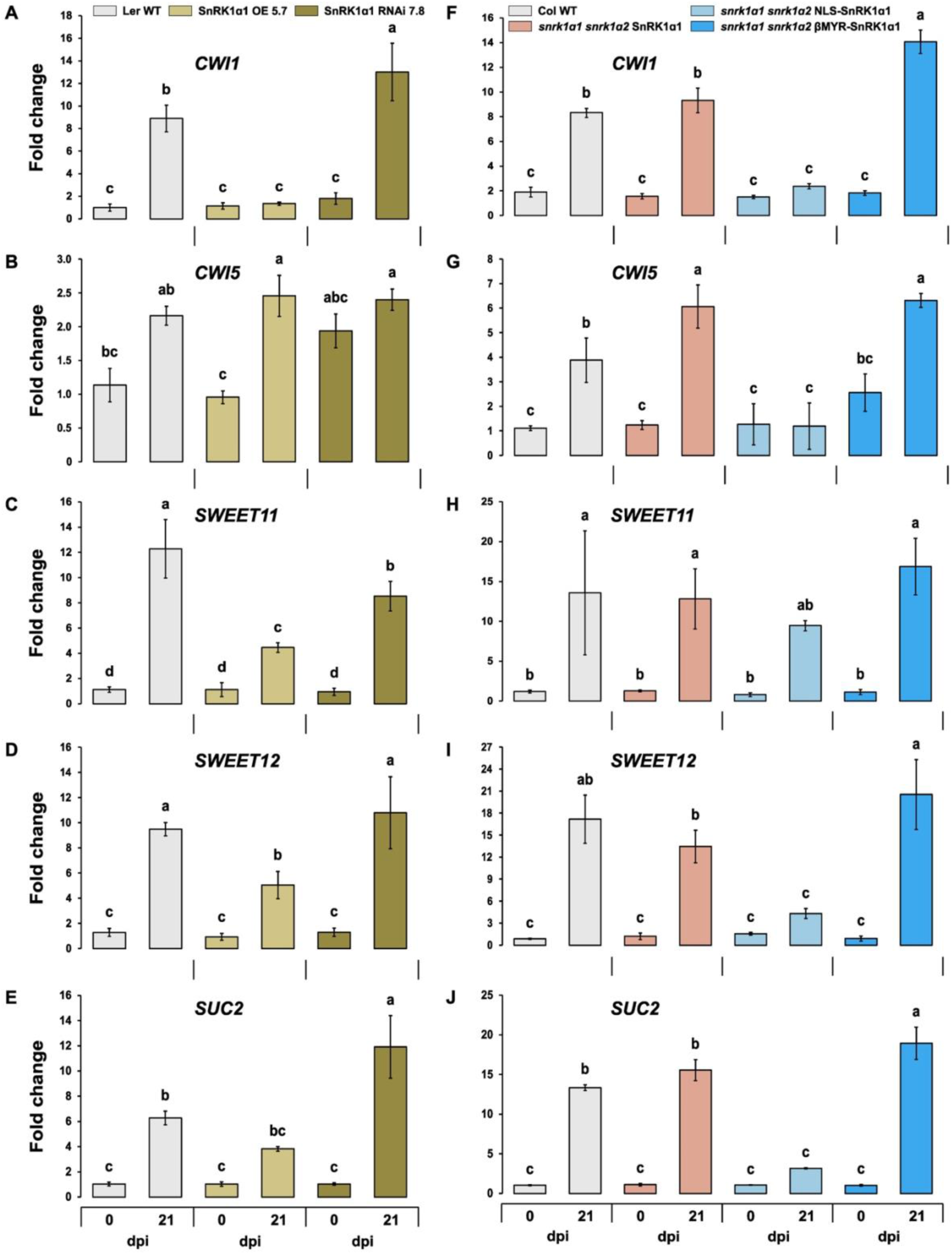
Increased nuclear SnRK1 activity limits the induction of cell wall invertases (*CWI1* & *CWI5*) and sugar transporters (*SWEET11*, *SWEET12* & *SUC2*) in roots during clubroot infection. (A-B, F-G) Cell wall invertase (*CWI1* & *CWI5*) expression, (C-D, H-I) expression of *SUGARS WILL EVENTUALLY BE EXPORTED TANSPORTERS* (*SWEET11* & *SWEET12*) and (E, J) *SUCROSE TRANSPORTER2 (SUC2)* were measured in infected roots of different SnRK1α1 modified-lines from hydroponically grown *A. thaliana* at 0 and 21 dpi using qRT-PCR. Data are the mean ± SD of three independent biological replicates, a total of 60–80 plants. Relative expression was normalized with the expression of *AtTUB* and calculated against 0 dpi. Error bars represent the standard error of the means of the three replicates. Different letters represent statistically significant differences across different genotypes (p < 0.005; two-way analysis of variance).

In summary, these analyses revealed dynamic changes in *CWI1* and *CWI5* expression, consistent with CWINV activities. While clubroot infection induces sink-related proteins, increased nuclear SnRK1 activity counteracts this response by transcriptionally repressing their expression, most prominently that of CWI1, thereby reducing sucrose cleavage and limiting carbon supply to developing galls.

### SnRK1 also affects sugar transporter gene expression during clubroot infection

SUGARS WILL EVENTUALLY BE EXPORTER TRANSPORTERS (SWEETs) play a crucial role in the translocation of sucrose and hexose to the phloem apoplasm, facilitating their subsequent import in companion cells and thus phloem loading and transport to sink tissues such as (infected) roots. They also contribute to phloem unloading in sink tissue and the importance of SWEETs in fueling clubroot development was demonstrated using *Arabidopsis sweet11* single and *sweet11 sweet12* double mutants, where disease progression was significantly delayed (Walerowski *et al*. 2018).

To investigate the influence of SnRK1 on *SWEET* (specifically *SWEET11* and *SWEET12*) expression, we analyzed their expression patterns in *P. brassicae*-infected roots across the different genotypes and time points. *SWEET11* and *SWEET12* expression increased over time in infected roots in all genotypes. However, this increase was significantly more pronounced in wild-type plants and susceptible lines, including SnRK1α1 RNAi, *snrk1α1*, and βMYR-SnRK1α1 lines, compared to SnRK1α1 overexpression (OE) and NLS-SnRK1α1 lines in both the Ler-0 and Col-0 ecotypes (Figure 7F-I, Suppl Figure S10, S11, S12).

SUCROSE TRANSPORTER 2 (SUC2) is a plasma membrane-localized H⁺/sucrose-symporter in companion cells that actively mediates sucrose loading from the apoplast into the phloem, enabling the efficient long-distance transport of photoassimilates from source leaves to sink tissues such as (infected) roots. By maintaining sucrose translocation, SUC2 is essential for carbon allocation, plant growth, and metabolic support of pathogen-infected organ development. Interestingly, SUC2 is also highly expressed in sink tissues, including roots, where it contributes to sucrose re-loading by retrieving sucrose that escapes from the phloem translocation stream into surrounding tissues (Gould *et al*. 2012). Although a possible role of SUC2 in more directly mediating sucrose unloading in sink tissues has been suggested, experimental evidence is still lacking. We analyzed *SUC2* transcriptional dynamics in *P. brassicae*-infected roots across different genotypes and time points. In all genotypes, *SUC2* transcript levels gradually increased during infection. However, the induction was markedly higher in the wild-type and susceptible SnRK1α1 RNAi, *snrk1α1*, and βMYR-SnRK1α lines, compared to the SnRK1α1 overexpression (OE) and NLS-SnRK1α1 lines. This pattern was consistent across both the Ler-0 and Col-0 ecotypes, except for Col SnRK1α1 OE 1.3, which showed increased expression at later time points (Figure 7E, 7J, Suppl Figure S13).

These findings suggest that SnRK1 not only modulates sugar dynamics in infected roots but may also (directly or indirectly) regulate the expression of key sugar transporters during clubroot infection, further underscoring the importance of SnRK1 in shaping host-pathogen interactions and nutrient allocation during disease progression at least in part through (nuclear) transcriptional reprogramming.

### The *P. brassicae* effector PBZF1 limits SnRK1α1 nuclear localization

The *P. brassicae* Arg-x-Leu-Arg motif and zinc finger-containing effector PBZF1 was reported to directly target SnRK1α1 (Chen *et al*. 2021). Cytoplasmic effectors with an RxLR motif are secreted by a range of pathogens to manipulate host immunity and enhance virulence (Combier, Evangelisti, Piron, Schornack & Mestre 2022; Wang, McLellan, Boevink & Birch 2023). With remarkably varied modes of action, RxLR effectors can target a wide range of host proteins, inhibiting enzyme activities, disrupting protein complexes, and destabilizing or relocating their targets (Wang *et al*. 2023). Following the reported interaction and inhibitory effect of PBZF1 on SnRK1α1 target gene activation (Chen *et al*. 2021), we more directly examined the impact of PBZF1 on SnRK1α1 activity using transient expression in *Arabidopsis* leaf mesophyll protoplasts and the established *DARK INDUCED6 (DIN6)/ASPARAGINE SYNTHASE1* promoter-luciferase (LUC) reporter as a read-out of nuclear SnRK1 activity (Baena-González *et al*. 2007; Dietrich *et al*. 2011). Co-expression of (HA-tagged) PBZF1-HA with SnRK1α1-HA (confirmed using immunoblotting) significantly reduced the ability of SnRK1α1 to activate the *DIN6* promoter, confirming PBZF’s inhibitory effector function (Figure 8A, B). Interestingly, PBZF1 was less effective at inhibiting *DIN6-LUC* activation by the NLS-SnRK1α1 protein carrying a strong NLS (Figure 8C). To further investigate the mechanism by which the effector affects SnRK1α1 function, we examined the subcellular localization of both proteins in *Arabidopsis* leaf mesophyll protoplasts using GFP-tagged constructs and confocal fluorescence microscopy. Under control conditions, SnRK1α1 localized both in the nucleus and cytosol of leaf mesophyll cells, as previously reported and consistent with its dynamic nuclear translocation for target gene regulation (Ramon *et al*. 2019). PBZF1, however, uniquely localized in the cytosol in our assays, consistent with the reported cytosolic interaction (Chen *et al*. 2021). Remarkably, upon co-expression, the nuclear SnRK1α1-GFP signal appeared to decrease significantly, indicating that PBZF1 restricts nuclear translocation of the kinase (Figure 8D). Quantification of the nuclear co-localization and the nuclear-to-cytoplasmic GFP-ratio of SnRK1α1 in the presence or absence of PBZF1, based on confocal imaging of leaf mesophyll protoplasts using Fiji (ImageJ), strongly supports this observation (Suppl Figure S14). To corroborate this effect in a system that more closely reflects the natural infection site of *P. brassicae*, we set up root protoplast isolation from hydroponically grown plants. While the transfection efficiency of root protoplasts was still too low for promoter-LUC reporter assays (requiring further optimization), fluorescence microscopy of GFP-positive cells clearly confirmed nuclear exclusion of SnRK1α1 in the presence of PBZF1. As a control, the reported abscisic acid (ABA)-triggered nuclear export of SnRK1α1 (Belda-Palazón, Costa, Beeckman, Rolland & Baena-González 2022) was confirmed in this experimental system (Figure 8E). In conclusion, our findings indicate that PBZF1 blocks SnRK1 target gene reprogramming (required to mount an effective defense response) by preventing its nuclear translocation.

**Figure 8.**
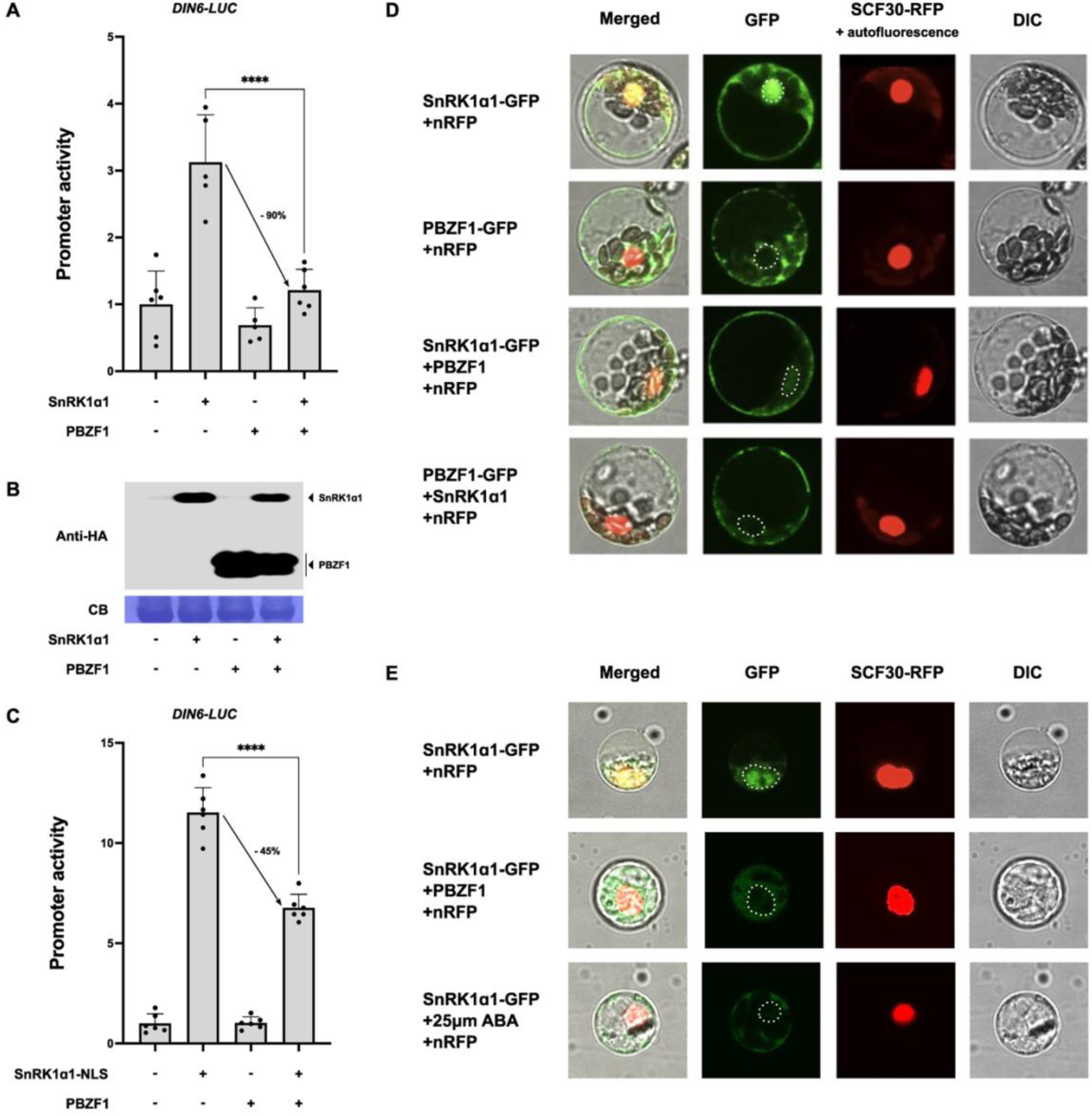
The *P. brassicae* effector PBZF1 inhibits SnRK1α1 activity by preventing its nuclear translocation. *DIN6* promoter activity was measured following transient co-expression of *PBZF1* with either *SnRK1α1* (A) or *NLS-SnRK1α1* (C) in Arabidopsis (Col-0) leaf mesophyll protoplasts. Transfected protoplasts were incubated for 6h at RT under continuous light conditions (75-100 µE). An empty vector was used as a negative control and a prUBQ10::GUS reporter construct served as an internal control. Values are averages with standard deviations, n=6. One-way ANOVA statistical analysis was performed, **** P<0.0001. (B) Immunoblot analysis with anti-HA antibodies to assess gene expression of *SnRK1α1* and *PBZF1*. Ribulose bisphosphate carboxylase small chain (RBCS) staining with Coomassie Brilliant Blue R-250 served as a protein loading control. (C) Subcellular localization of SnRK1α1-GFP and PBZF1-GFP in Arabidopsis leaf mesophyll protoplasts upon co-expression of either *SnRK1α1* or *PBZF1*. Subcellular localization was visualized 14h after transfection using confocal fluorescence microscopy (FluoViewTM FV1000 confocal system). An SCF30-RFP (nRFP) construct was co-expressed and served as a red fluorescent nuclear marker. (D) Subcellular localization of SnRK1α1-GFP in Arabidopsis root protoplasts upon co-expression of *PBZF1*. Subcellular localization was visualized 20h after transfection using confocal fluorescence microscopy.

## Discussion

In this study, we investigated the role of SnRK1 in *Arabidopsis* defense against *P. brassicae* infection and clubroot disease. To overcome limitations associated with soil-based assays, including heterogeneous pathogen exposure and restricted access to intact root tissues, we established a hydroponic infection system that enables more uniform inoculation and precise monitoring of disease progression (Figure 3A). Beyond facilitating phenotyping, this platform will also enable more effective use of reporter lines and single-cell and spatial omics approaches. Using complementary soil- and hydroponics-based assays, we show that enhanced SnRK1 activity delays clubroot development, whereas reduced SnRK1α1 function exacerbates disease severity across genetic backgrounds (Figure 1 and 3, Suppl Figure S1-3). These findings are consistent with, and extend, independent reports identifying SnRK1α1 as a positive regulator of resistance against clubroot (Chen *et al*. 2021), and support a conserved role for SnRK1 in restricting pathogen proliferation.

Clubroot disease is characterized by extensive metabolic reprogramming of infected roots, particularly in carbohydrate allocation, which underpins gall formation and pathogen growth. Elevated levels of sucrose, trehalose, and starch in infected tissues have been reported across multiple host species and experimental systems, reflecting a redirection of carbon toward developing galls and the establishment of strong metabolic sinks (Keen & Williams 1969; Evans & Scholes 1995; Brodmann *et al*. 2002; Kong *et al*. 2022). Support for this sink-based model comes from both comparative and molecular studies: resistant *Brassica rapa* lines accumulate lower levels of soluble sugars than susceptible lines (Li *et al*. 2018), while in *Arabidopsis*, *P. brassicae* infection induces the expression of sugar transporters and carbohydrate metabolic genes, with transcript abundance correlating with sugar accumulation in infected roots (Siemens *et al*. 2006; Irani *et al*. 2018; Li *et al*. 2018; Walerowski *et al*. 2018; Zhang *et al*. 2019; Wang *et al*. 2019b; Ciaghi, Schwelm & Neuhauser 2019; Adhikary *et al*. 2022).

Based on SnRK1’s established role as a master regulator of cellular energy homeostasis, we hypothesized that SnRK1 functions as a metabolic gatekeeper, limiting carbon allocation to infected tissues during clubroot development. Our data support this hypothesis as reduced SnRK1α1 activity resulted in elevated hexose levels in infected roots, accompanied by increased expression of sugar transporters and cell wall invertase genes and enhanced invertase activity. Conversely, SnRK1α1 overexpression reduced hexose levels and attenuated the induction of sugar transporters and invertases following infection (Figure 5 and 7, Suppl Figure S5, S6, S8-S11). These effects were observed consistently across Col-0 and Ler-0 backgrounds.

Nuclear localization of SnRK1α1 is critical for transcriptional regulation of SnRK1 target genes and thus SnRK1-mediated transcriptional metabolic and defense responses. Consistently, enhanced nuclear SnRK1α1 localization effectively delayed clubroot disease development, while cytosolic retention of SnRK1α1 significantly exacerbated the phenotypes (Figure 2 and 3, Suppl Figure S4). This was associated with reduced and increased hexose levels and sugar transporter and invertase gene expression, respectively (Figure 6 and 7, Suppl Figure S7-9, S12). These findings support a model in which nuclear translocation of SnRK1α1 limits the establishment of pathogen-driven carbon sinks, also making this mechanism a putative target for pathogen effectors to insure carbon and energy supply.

Among the SnRK1-regulated processes, SUC2- and SWEET-mediated sucrose transport and CWI-mediated sucrose hydrolysis thus represent central nodes controlling source-to-sink carbon allocation. Elevated expression of the transporters during *P. brassicae* infection likely enhances carbon flux toward developing galls. The increased expression of *SUC2*, *SWEET* and *CWI* genes in SnRK1α1 RNAi lines, together with their repression in overexpression lines, indicates that SnRK1 directly restrains pathogen-driven carbon and energy supply. Functional evidence supports this model: impaired sucrose transport in the *sweet11 sweet12* double mutant reduces disease severity, and suppression of cell wall invertase activity through root-specific invertase inhibitors markedly attenuates gall development (Siemens *et al*. 2011; Walerowski *et al*. 2018). These processes are further reinforced by elevated cytokinin levels during infection, which enhance sugar transporter and invertase gene expression and thereby strengthen metabolic sinks (Ehness & Roitsch 1997; Devos *et al*. 2006; Siemens *et al*. 2006). Collectively, these results position SnRK1 upstream of key metabolic nodes, acting to restrict pathogen-induced carbon diversion and limit clubroot development.

The role of SnRK1 in controlling carbon metabolism extends beyond *Arabidopsis*, highlighting a conserved function across plant species and its potential as a target for crop improvement. In potato and sweet potato, SnRK1 OE enhances starch accumulation while reducing glucose levels, reflecting its impact on sugar partitioning and invertase activity (McKibbin *et al*. 2006; Lin *et al*. 2015; Ren, He, Zhao, Zhai & Liu 2019). Similarly, maize ZmSnRK1 OE in *Arabidopsis* increases sucrose content and induces significant physiological changes (Wang *et al*. 2019a), whereas SnRK1-silenced cucumber lines display elevated soluble sugar levels compared to OE lines (Huang *et al*. 2023). In various fruit crops, SnRK1 OE boosts sucrose accumulation and modulates key sucrose metabolism genes, with effects varying according to source and sink tissues: leaves primarily export sugars, whereas roots and fruits act as strong importers (Wang, Peng, Li, Yang & Li 2012; Yu, Peng, Xiao, Wang & Luo 2018; Luo, Peng, Zhang, Xiao & Zhang 2020). These findings suggest that SnRK1-mediated regulation of carbon allocation is broadly conserved and could be leveraged to improve crop resilience and productivity.

Mechanistically, SnRK1 can also directly target sugar metabolic enzymes. Sucrose-P synthase (SPS), a key rate-limiting enzyme in sucrose biosynthesis, is one of the first identified direct phosphorylation targets, inactivated by SnRK1 (Sugden, Donaghy, Halford & Hardie 1999). Interestingly, cytosolic invertase CINV1 has been identified as a potential SnRK1 interactor and phosphorylation target in *Arabidopsis* (Van Leene *et al*. 2022), while in potato, invertase activity is finetuned by a complex containing invertase, its inhibitor, and SnRK1 (Lin *et al*. 2015). Such interactions highlight SnRK1’s capacity to coordinate both transcriptional and post-translational control of carbohydrate allocation. Notably, soluble sugar accumulation can feedback to inhibit SnRK1 activity, implying that pathogen-induced sugar buildup during *P. brassicae* infection may dampen endogenous SnRK1 signaling, thereby facilitating pathogen proliferation. Collectively, these results underscore a conserved and multifaceted role for SnRK1 in regulating sugar metabolism, linking energy homeostasis to defense and offering a potential strategy for engineering crops with improved resistance to pathogen-induced metabolic reprogramming.

Beyond metabolic regulation, SnRK1 plays a pivotal role in plant immunity, mediating resistance against a broad spectrum of pathogens in different economically important crop species. In rice, OsSnRK1 isoforms positively regulate basal resistance against fungal blast and bacterial blight, contributing to broad-spectrum defense against necrotrophic and (hemi)biotrophic pathogens (Kim *et al*. 2015; Filipe *et al*. 2018). Overexpression of OsSnRK1α enhances resistance to rice blast by positive regulation of the salicylic acid (SA) pathway and jasmonate-mediated defense responses (Cao *et al*. 2024). In barley, SnRK1 phosphorylates the WRKY3 transcription factor, a repressor of basal immunity, thereby enhancing resistance to powdery mildew (Han *et al*. 2020). Similarly, in bread wheat, TaSnRK1α overexpression confers tolerance to *Fusarium graminearum*, whereas RNAi lines are more susceptible (Jiang *et al*. 2020). In strawberries, SnRK1 enhances resistance to *Botrytis cinerea* through SA signaling, potentially integrating metabolic cues with immune activation (Luo *et al*. 2024).

During infection, *P. brassicae* manipulates host hormone signaling to promote gall formation, primarily by enhancing (growth-stimulating) cytokinin and auxin biosynthesis while suppressing SA-dependent defense (Siemens *et al*. 2006; Jahn *et al*. 2013; Ludwig-Müller 2014; Ludwig-Müller *et al*. 2015; Lemarié *et al*. 2015; Lovelock *et al*. 2016; Malinowski *et al*. 2016; Xu *et al*. 2018). Notably, in *Arabidopsis*, SnRK1 was found to have a dual role in SA-mediated defense: it positively regulates NPR1-dependent immunity while fine-tuning alternative SA-independent pathways to prevent excessive activation under stress (Wang et al. 2019; Jie et al. 2025). Moreover, SA itself can activate SnRK1 signaling, promoting phosphorylation of NPR1 and reinforcing effective defense responses (Chen *et al*. 2025). These observations suggest that SnRK1 integrates metabolic status with immune signaling across plant species, allowing plants to balance growth, carbon allocation, and defense in response to pathogen attack.

Early host responses include PAMP-triggered immunity (PTI), characterized by reactive oxygen species bursts and callose deposition, alongside activation of salicylic acid (SA)–responsive defense pathways. Transcriptomic analyses of resistant genotypes reveal ETI-like gene expression patterns, including effector receptor homologs and SA-responsive defense genes, suggesting that recognition of pathogen effectors may contribute to early immune signaling (Chen *et al*. 2021; Adhikary *et al*. 2022; Wang *et al*. 2022; Oh, Park, Lee, Shim & Oh 2024). SnRK1 likely participates in these early defenses by regulating energy allocation to support immune responses and by modulating transcriptional programs associated with both PTI and ETI-like pathways. Importantly, effectors such as PBZF1 are expressed early during infection, including in resting spores and primary infection stages. As PBZF1 interacts with SnRK1α1 specifically in the cytosol and not in the nucleus (Chen *et al*. 2021), we hypothesized that it interferes with nuclear translocation to suppress transcriptional defense programs. Our GFP assays confirmed this mechanism in both leaf mesophyll and root protoplasts (Figure 8D–E), providing direct evidence that effector-mediated modulation of SnRK1 subcellular localization contributes to early suppression of host immunity. Collectively, these results support a model in which SnRK1 activity from the outset of infection is critical for limiting pathogen growth. Effector-mediated suppression of SnRK1 has also been reported in other plant–pathogen systems, including wheat–stripe rust (*Pst4121*) and rice fungal pathogens (Qian *et al*. 2025). Also, the SnRK1 activating kinases SnAK1 and SnAK2 were first identified as geminivirus replicase proteins Rep/AL1-interacting kinases (GRIK1/2), that highly accumulate in infected tissue, most likely to limit the virus’ repressive effect on SnRK1 activity to enable its proliferation (Kong & Hanley-Bowdoin 2002; Shen & Hanley-Bowdoin 2006).

Indeed, another notable feature of clubroot pathology is its effect on host growth and development, including manipulation of the cell cycle machinery and root vascular differentiation. The pathogen’s induction of cell cycle genes is consistent with cytokinin/auxin-driven activation of cyclin-dependent kinases (Del Pozo, Lopez-Matas, Ramirez-Parra & Gutierrez 2005; Olszak *et al*. 2019). SnRK1 may impose a metabolic checkpoint for growth, for exaple by phosphorylating KRP inhibitors, antagonizing the TOR kinase, or directly modulating the E2F TF, thereby limiting resource-intensive gall proliferation (Xiong *et al*. 2013; Guérinier *et al*. 2013; Nukarinen *et al*. 2016; Broeckx *et al*. 2016; Son *et al*. 2023). Through such mechanisms, in an intricate nutrient- and stress-responsive network with TOR kinase, SnRK1 can balance growth-defense trade-offs (Crepin & Rolland 2019). By coordinating energy homeostasis and sugar allocation, SnRK1 could counteract the pathogen-driven shift toward enhanced phloem differentiation and suppressed xylogenesis, which establishes a strong metabolic sink for pathogen proliferation (Malinowski, Smith, Fleming, Scholes & Rolfe 2012; Malinowski, Truman & Blicharz 2019; Walerowski *et al*. 2018). While a direct molecular link between SnRK1 and vascular development remains to be identified, our findings indicate that SnRK1 also gatekeeps growth and developmental processes to restrict nutrient flow to and proliferation of infected tissues.

In summary, our study identifies SnRK1 as a central integrator of metabolic, immune, and developmental signaling during *Plasmodiophora brassicae* infection. Restricting carbon allocation to infected tissues, at least in part through regulation of sugar transporter and invertase activity by nuclear transcriptional reprogramming, SnRK1 activity acts as a constraint on pathogen-induced sink establishment and gall development. Our findings further demonstrate that this regulatory node is actively targeted by the *P. brassicae* effector *PBZF1*, that interferes with SnRK1α1 nuclear translocation, thereby suppressing host defense programs. Together with evidence from other plant–pathogen systems, these results support a model in which pathogens exploit SnRK1’s central position to uncouple host metabolism from immunity and growth control. While broad activation of SnRK1 likely imposes growth penalties, resolving the specific downstream branches and spatial regulation of SnRK1 signaling offers a promising strategy for a better balancing of growth and defense expenses and a more durable, quantitative resistance.

## Methods and Materials

### Plant Material and Growth Conditions

The different lines of *Arabidopsis thaliana* used in this study were in the Col-0 or Ler-0 background. The Ler SnRK1α1 OE, Ler SnRK1α1 RNAi (Baena-González *et al*. 2007), Col OE lines, *snrk1α1* T-DNA, *snrk1α1 snrk1α2* SnRK1α1, *snrk1α1 snrk1α2* NLS*-*SnRK1α1, *snrk1α1 snrk1α2* βMYR-SnRK1α1 mutants and transgenic lines (Ramon *et al*. 2019) have been described previously. Col OE lines were produced using a pCB302-based binary vector with 35SC4PPDK promoter (35S enhancer and maize C4PPDK basal promoter) and nopaline synthase (NOS) terminator (Sheen 1996). The seeds were sterilized by vapor and then stratified for three days at 4°C. Plants were typically grown in plant growth chambers under a 12-hour light/dark cycle with cool white, fluorescent light at 21°C. For soil disease bioassays, plants were grown in a soil-vermiculite mixture of 3:1 (w:w) in a plant growth chamber in a 12-h light/12-h dark diurnal cycle at 21°C with 100 μmol m^-2^ s^-1^ cool white, fluorescent light.

### Simplified hydroponic system for *P. brassicae* disease bioassays

A simplified hydroponic system, adapted from Conn and colleagues with significant modifications (Conn *et al*. 2013), was employed for the *P. brassicae* disease bioassays. Central apertures were bored through black microcentrifuge tube lids. These lids were then filled with approximately 250-300 μL of ½ MS medium with agar, allowing solidification for 15 minutes. Subsequently, the agar-containing lids were placed into custom-designed nine or 24-well black plexi holders tailored to fit standard tip boxes. This setup facilitated their function as platforms for supporting seeds, seedlings, or plants. A modified ½ Hoagland medium was used as a growing medium for the *Arabidopsis* plants (Conn *et al*. 2013). The tip boxes were filled with approximately 240 – 260 mL ½ Hoagland medium. Following this, three to five stratified and sterilized seeds were planted in each agar-filled 24-well holder for germination. The lids containing the germinated seeds were transferred to 9-well holders between 7 and 10 days after sowing to initiate *P. brassicae* infection. At this stage, only one seedling was retained in each lid and well. The tip boxes with the seedlings were typically incubated in a plant growth chamber in a 12-h light/12-h dark diurnal cycle at 21°C with 100 μmol m^-2^ s^-1^ cool white, fluorescent light. The Hoagland medium was replenished every 2 to 3 days.

### *Plasmodiophora brassicae* disease bioassays and disease scoring

For all the experiments, the *P. brassicae* e3 single spore isolate was used for soil and hydroponics disease bioassays (Fähling, Graf & Siemens 2003). The pathogen was cultured in *B. rapa* plants to promote its growth and reproduction. Pathogen spores were isolated following previously established protocols (Jahn *et al*. 2013). At 14 days after sowing, each seedling was inoculated with 1 mL of e3 single spore suspension (10^7^ spores/mL in 50 mM KH_2_PO_4_, pH 5.5). For the *P. brassicae* hydroponic assay, *A. thaliana* plants were grown following the instructions provided in the simplified hydroponic system for the *P. brassicae* disease bioassays. At 14 days after sowing, tip boxes were infected with 250 to 300 µL of *P. brassicae* e3 single spore isolate (10^9^ spores/mL in 50 mM KH_2_PO_4_, pH 5.5), and the solution was mixed well using a pipette.

Disease severity was assessed 26-28 days after inoculation by removing the roots from the soil or hydroponic system, and symptom severity was scored according to (Siemens, Nagel, Ludwig-Müller & Sacristán 2002) categorizing plants into five distinct classes: Class 0 plants devoid of any disease symptoms; Class 1 plants exhibiting minor swellings at the minor and/or secondary roots while maintaining typical root structure; Class 2 plants with visibly thickened primary roots, reduced fine roots and lateral roots, and further thickening of affected roots; Class 3 plants were characterized with significantly reduced root systems displaying clearly visible galls on primary and secondary roots, with fine roots no longer discernible and occasional gall development on the hypocotyl; Class 4 plants with roots primarily composed of a single sizable brownish gall. Based on this classification, a Disease Index (DI) was calculated according to the following formula:

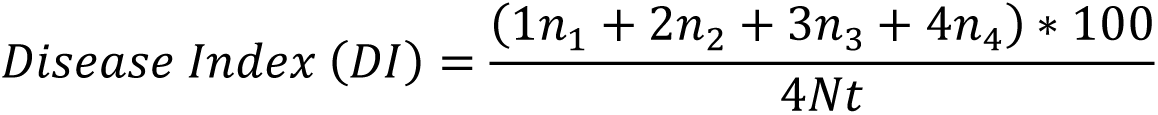

with n1 to n4 = number of plants in the different disease classes and Nt = Number of all tested plants.

Also, the shoot index (SI) was calculated as the ratio of the fresh weight of infected plants to that of non-infected plants.

### Soluble sugar quantification and invertase activity determination

After infection with *P. brassicae*, we collected non-infected and infected root samples from hydroponically grown *Arabidopsis* plants at different time points (0 to 28 dpi days post-infection). For each sample at each time point, 15 to 18 non-infected/infected root tissues were sampled for one replicate, and the assay was quantified using three biological replicates. The extraction of soluble sugars and invertase activity (cell wall and vacuolar invertase) from plant root tissues was done according to (Versluys, Toksoy Öner & Van den Ende 2022). Ground root powder samples (100 mg) were boiled at 100°C in ddH_2_O for 10 minutes. Following centrifugation at 15,000 g for 10 minutes, 200 µL of the resulting supernatant was transferred to a Dowex® column containing H^+^ and Ac^−^ resins. The column underwent six washes with 200 µl ddH_2_O, and the entire eluate was collected. Subsequently, the collected volume was centrifuged for 5 minutes at 15,000 g and analyzed to determine the concentrations of soluble sugars Glc, Fru, and Suc utilizing HPAEC-IPAD (High-Performance Anion-Exchange Chromatography with Integrated Pulsed Amperometric Detection).

For cell wall (CWINV) and vacuolar invertase activity (VINV), Ground root tissue samples (100 mg) were homogenized in 500 µl ice-cold extraction buffer consisting of 50 mM NaOAc (pH 5.0), 10 mM NaHSO_3_, 2 mM β-mercaptoethanol, 0.1% polyclar, and 100 µM phenylmethylsulfonyl fluoride. Following extraction on ice using a micro pestle, the samples underwent centrifugation for 20 minutes at 20,000 g and 4°C. The resulting pellets contained the CWINV fraction, while VINV activity was assessed in the supernatants. Pellets were subjected to several washes with 50 mM NaOAc buffer (pH 5.0), followed by centrifugation for 10 minutes at 20,000 g and 4°C and resuspension using a micro pestle. In the final resuspension step, 500 µl buffer containing 50 mM sucrose was added to initiate the reaction. After 30 minutes of incubation at 30°C with shaking at 600 rpm, the samples were heated to 95°C for 5 minutes. VINV was further purified from the supernatant fraction through two rounds of ammonium sulfate precipitation (80%) on ice, followed by centrifugation at 4°C for 5 minutes at 20,000 g. The resulting pellet was resuspended in 50 mM NaAc buffer, and the reactions were incubated at 30°C for 30 minutes in NaOAc buffer (pH 5.0) containing 100 mM sucrose. Control samples were withdrawn at 0 minutes and immediately heated. Following a final centrifugation for 5 minutes at 15,000 g, invertase activity was assessed by HPAEC-IPAD quantifying the increase in fructose. Enzyme activity was expressed as the production of fructose from sucrose, measured in nmol min^–1^ g^–1^ FW.

### RNA extraction and qRT-PCR

RNA extraction from non-infected and infected root tissues was performed using TRIzol^TM^ Reagent (Bioline, London, UK) following the manufacturer’s protocol. The isolated RNA was then reverse-transcribed into cDNA using the SensiFAST™ cDNA synthesis kit (Bioline). Quantitative real-time PCR (qRT-PCR) was performed in a 96-well plate using the StepOne™ Real-Time PCR System (Thermo Fisher Scientific, Waltham, MA, USA). PowerUp™ SYBR™ Green Master Mix and specific primers (Table S1) were utilized for qRT-PCR amplification (Thermo Fisher Scientific). Each reaction mixture contained 10 nanograms of cDNA in a total volume of 10 μL, comprising 5 μL of PowerUp SYBR Green master mix, 0.2 μl of each primer, and 2.6 μl of H_2_O. Thermal cycling conditions consisted of an initial denaturation step at 95°C for 2 minutes, followed by 40 cycles of denaturation at 95°C for 3 seconds and annealing/extension at 60°C for 30 seconds. Normalization of marker gene expression was performed relative to the expression of the UBQ10 gene, selected for its demonstrated stability across various tissues and metabolic stress conditions (Czechowski, Stitt, Altmann, Udvardi & Scheible 2005). All reactions were performed in triplicate, and the 2^-ΔΔCT^ method was applied to calculate the relative expression.

### In planta quantification of *Plasmodiophora brassicae*

Healthy and infected root tissues were collected from *Arabidopsis* plants at various time points post-inoculation. Total DNA was extracted from approximately 100 mg of fresh root tissue using a DNA extraction buffer (1.4 M NaCl, 20 mM EDTA, 100 mM Tris-HCl pH 8.0, 3% w/v CTAB, and 1% β-mercaptoethanol). The tissue was homogenized in the extraction buffer and incubated at 65°C for 30–60 minutes. The lysate was then subjected to chloroform:isoamyl alcohol (24:1) separation, followed by centrifugation at 12,000 rpm for 10 minutes to isolate the aqueous phase. DNA was precipitated from the aqueous phase by adding 0.7 volumes of isopropanol and incubating at -20°C for 30 minutes. The DNA pellet was washed with 70% ethanol, air-dried at room temperature, and resuspended in TE buffer or nuclease-free water. To quantify *P. brassicae* DNA, we used *P. brassicae*-specific Internal Transcribed Spacer (ITS) regions within the ribosomal DNA and PbAction as a reference gene for *P. brassicae* (Sundelin *et al*. 2010). The primers used for the quantification are listed in Table S1. All reactions were performed in triplicate, and the 2^-ΔΔCT^ method was applied to calculate the relative expression.

### Transient expression in leaf mesophyll and root protoplasts

Isolation and PEG/Ca^2+^ transfection of *Arabidopsis* leaf mesophyll protoplasts was performed as described by (Yoo, Cho & Sheen 2007). For root protoplast isolation, cells were harvested from 10-day-old *A. thaliana* seedling roots. Leaf mesophyll protoplasts were transfected with CsCl-gradient purified plasmid DNA and incubated in dim light for 6h (LUC/GUS assay and immunoblot analyses) or 14 h (subcellular localization assay). For transfection of root protoplasts, 250 µL PEG was added to 250 µL protoplasts and 50 µg plasmid DNA. The PEG/Ca^2+^ treatment was extended to 20 min (instead of 5 min), and root cells were incubated in the dark for 20h (subcellular localization assay). Protoplasts were harvested by centrifugation (200*g*; RT) using a swinging bucket rotor (model 5804; Eppendorf, Hamburg, Germany) and cell pellets were stored at -80°C or immediately analyzed (microscopy). The SnRK1α1 and NLS-SnRK1α1 constructs with HA and GFP tag were previously described (Baena-González *et al*. 2007; Ramon *et al*. 2019). The *PBZF1* coding sequence (PBCN_001987) was PCR-amplified from *P. brassicae* cDNA (Table S1) and inserted in the HBT95 expression vector with the 35SC4PPDK promoter (35S enhancer and maize C4PPDK basal promoter) and nopaline synthase (NOS) terminator in-frame with a double HA or GFP tag (Sheen 1996).

### LUC and GUS assay

The pr*DIN6::LUC* and pr*UBQ10::GUS* constructs were described previously (Baena-González *et al*. 2007). Luciferase (LUC) and β-glucuronidase (GUS) assays were used for measuring relative (*DIN6*) target promoter activity. Transfected protoplasts were lysed with 100 µL lysis buffer (25 mM Tris-phosphate pH 7.8, 2 mM DTT, 2 mM 1,2-diaminocyclohexane-N,N,N’,N’-tetraacetic acid, 10% glycerol, 1% Triton X-100; Promega). 100 μL LUC reagent (E1500 Kit; Promega, Madison, WI, USA) was added to 20 µL of cell lysate, and luminescence was measured using a Lumat LB9507 tube luminometer (Berthold Technologies, Bad Wildbad, Germany). A pr*UBQ10::GUS* reporter was co-transfected as an internal control to correct for differential transfection efficiency and pipetting errors. 5 µL cell lysate was added to 45 µL of 10 mM 4-methylumbelliferyl β-D-glucuronide solution (MUG, M-9130; Sigma-Aldrich, St Louis, MO, USA) and incubated for 1 h at 37°C. The reaction was stopped by adding 220 µL of 0.2 M Na_2_CO_3_ and fluorescence was measured using a Glomax® multidetection system (Promega).

### Subcellular localization assay

Subcellular localization of transiently overexpressed proteins was examined at 40x magnification (488 nm) using the FluoViewTM FV1000 confocal system (Olympus). *Arabidopsis* leaf mesophyll and root protoplasts were transfected with plasmid DNA of eGFP-tagged constructs and incubated for 14 h and 20 h, respectively, before visualization.

### Immunoblot analyses

Protein levels were confirmed using immunoblot analysis. Before SDS-PAGE (Polyacrylamide Gel Electrophoresis), 20 µL loading buffer (120 mM Tris–HCl at pH 6.8, 5.4 M urea, 20% [v/v] glycerol, 4% [w/v] SDS, 5% [v/v] β-mercaptoethanol, and 0.5% [v/v] bromophenol blue) was added to the samples, which were then heated for 5 min at 95°C (VWR® Analog Heat Block). Proteins were separated on a 1.5 mm, 10% acrylamide SDS-PAGE gel in Tris-Gly running buffer (25 mM Tris, 192 mM Gly, and 0.1% [w/v] SDS at pH 8.5). Electrophoresis was performed with an initial stacking phase at 60 V for 15 min, followed by 15 min at 110 V, and a final separation phase at 160 V for 1 h. Separated proteins were transferred to a polyvinylidene fluoride (PVDF) membrane (Immobilon-P; Millipore, Burlington, MA, USA) using a wet blot system (MiniPROTEANTetra Cell; Bio-Rad, Hercules, CA, USA) in Tris-Gly buffer containing 10% (v/v) methanol at 300 mA for 2 h. The membrane was subsequently blocked for 1 h with 5% skimmed milk and incubated overnight at 4°C with horseradish peroxidase (HRP)-conjugated antibody (HRP-conjugated anti-HA antibody, 1:1000 [50 mg ml^-1^], catalog. no. 12013819001, Roche, Basel, Switzerland). The membrane was washed three times with TBST (500 mM NaCl, 13.5 mM Tris-HCl pH 7.5, and 0.05% Tween 20) and incubated for 2 min in Supersignal West Pico Plus Chemiluminescent substrate (catalog. no. 34577; Thermo Fisher Scientific). Antibody-bound proteins were visualized using a luminescent image analyzer (LAS-4000 mini; Fujifilm). Ribulose bisphosphate carboxylase small chain (RBCS) staining with Coomassie Brilliant Blue R-250 served as a protein loading control.

### Statistical Analyses

Statistical analyses were performed using GraphPad Prism9 and the *aov* function in the R statistical software, incorporating the Tukey post hoc comparisons and the multcomp function for compact letter display. Disease scoring was analyzed using one-way ANOVA, as this involved a single factor. For soluble sugar content, INV activity, and gene expression, two-way ANOVA was applied to assess the effects of genotype and time-point across two experimental conditions. The two-way ANOVA was conducted within each genotype to account for differences in response over time. A completely randomized design was employed, with three biological replicates, and each replicate consisted of a pool of at least 18 plants. Data were presented as mean ± standard error. Tukey’s post hoc test was used for multiple comparisons to assess significant differences between groups, with statistical significance set at p < 0.05.

## Acknowledgements

The authors are grateful to Jutta Ludwig-Müller for sharing e3 spore isolate and their advice. Hilde Verlinden, Anja Vandeperre, Maxime Versluys, and Timmy Reijnders provided excellent technical assistance and plant care. This research was funded by KU Leuven (C24/16/013 to F.R.) and FWO (SB/1S03125N to N.G.).

## Author Contributions

FR supervised the study. HV (disease assays, metabolite measurements, expression analyses), NG (effector studies), LV (initial observations with soil assays), and FR performed experiments and analyzed data. PVD, WVdE, and BDC provided feedback and advise as well as technical support throughout the project. HV, NG and FR wrote the manuscript. All authors read and approved the manuscript submitted for publication.

