## Supplementary data for "Nuclear SnRK1 activity delays clubroot development in *Arabidopsis* by reducing sink strength"

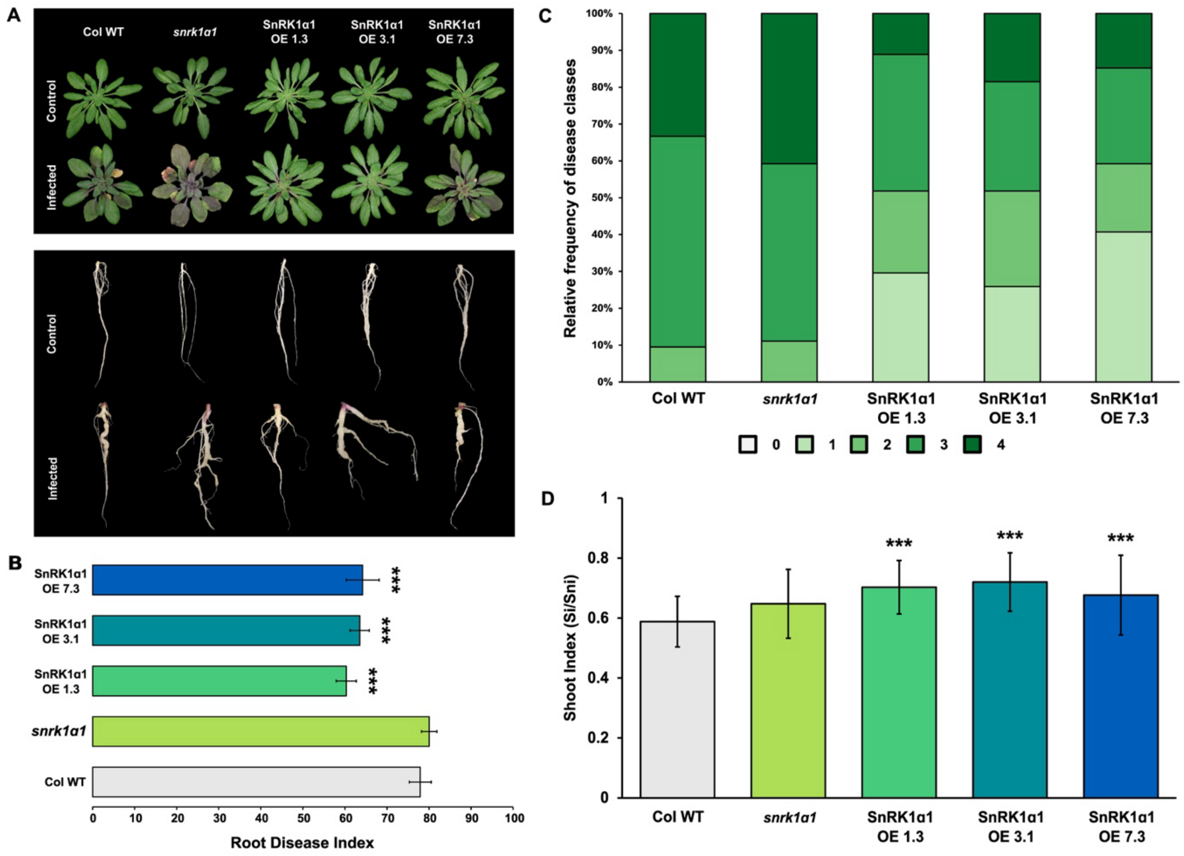


**Supplementary Figure 1. SnRK1α1 mediates resistance against clubroot disease development in soil-grown Col-0 background plants.** (A) Representative shoot and root phenotypes of soil-grown infected and uninfected (control) wild-type *A. thaliana* (Col-0 WT), three independent SnRK1α1 overexpression lines (OE 1.3, OE 3.1, and OE 7.3), and a single *snrk1α1* T-DNA line. (B) Root disease index (DI) quantifying overall disease severity for each genotype. (C) Relative frequency distribution of clubroot disease severity classes among the representative SnRK1α1 lines. Symptom severity was scored based on five distinct classes: Class 0, no visible symptoms; Class 1, minor swellings on minor and/or secondary roots while maintaining typical root structure; Class 2, visibly thickened primary roots, reduced fine roots and lateral roots; Class 3, significantly reduced root system with visible galls on primary and secondary roots, loss of fine roots, and occasional gall development on the hypocotyl; Class 4, roots primarily composed of a single sizeable brownish gall. (D) Relative shoot weight (shoot index, SI) was calculated for each line as the ratio of the fresh weight of infected plants to that of non-infected control plants (Si/Sni). Fourteen-day-old *A. thaliana* seedlings were inoculated with 1 mL of *P. brassicae* ‘e3’ single spore suspension ($10^9$ spores/mL in 50 mM $KH_2PO_4$, pH 5.5), and disease progress was evaluated at 28 days post-inoculation (dpi). Data are the mean ± SD of three independent biological replicates, a total of 60–80 plants. Error bars represent the standard error of the means of the three replicates. Asterisks indicate a significant difference compared with the wild-type (for *** $p < 0.001$; two-way analysis of variance).

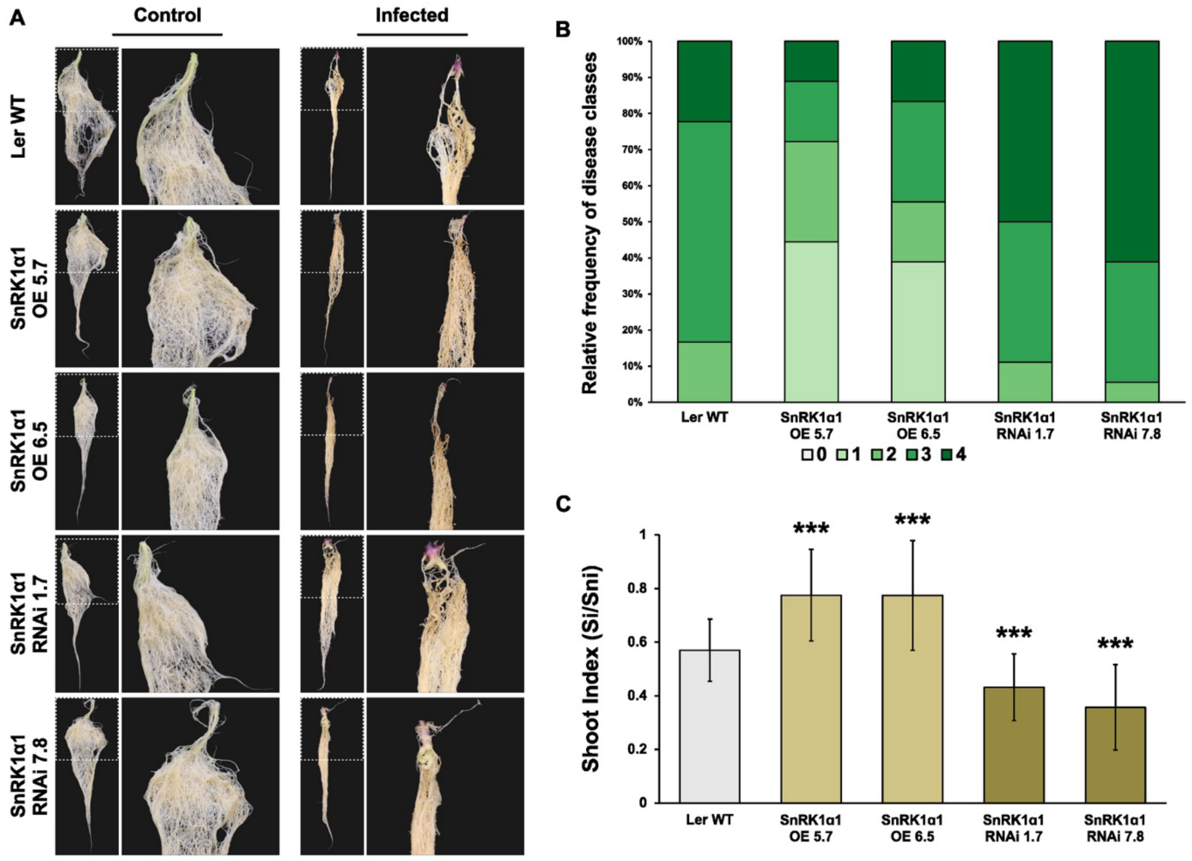


**Supplementary Figure 2. SnRK1α1 mediates resistance against clubroot disease development in Ler-0 background plants in a miniature hydroponics setup.** (A) Representative root phenotypes of infected and uninfected (control) wild-type *A. thaliana* (Ler-0 WT), two independent SnRK1α1 overexpression lines (OE 5.7 & OE 6.5), and two independent SnRK1α1 RNA interference lines (RNAi 1.7 & RNAi 7.8) grown in a miniature hydroponic system. (B) Relative frequency distribution of clubroot disease severity classes among the representative SnRK1α1 lines. Symptom severity was scored based on five distinct classes: Class 0, no visible symptoms; Class 1, minor swellings on minor and/or secondary roots while maintaining typical root structure; Class 2, visibly thickened primary roots, reduced fine roots and lateral roots; Class 3, significantly reduced root system with visible galls on primary and secondary roots, loss of fine roots, and occasional gall development on the hypocotyl; Class 4, roots primarily composed of a single sizeable brownish gall. (C) Relative shoot weight (shoot index, SI) was calculated for each line as the ratio of the fresh weight of infected plants to that of non-infected control plants (Si/Sni). Fourteen-day-old *A. thaliana* seedlings were inoculated with *P. brassicae* 'e3' single spore suspension ($10^9$ spores/mL in 50 mM $KH_2PO_4$, pH 5.5), and disease progress was evaluated at 28 days post-inoculation (dpi). Data are the mean ± SD of three independent biological replicates, a total of 60–80 plants. Error bars represent the standard error of the means of the three replicates. Asterisks indicate a significant difference compared with the wild-type (for *** $p < 0.001$; two-way analysis of variance).

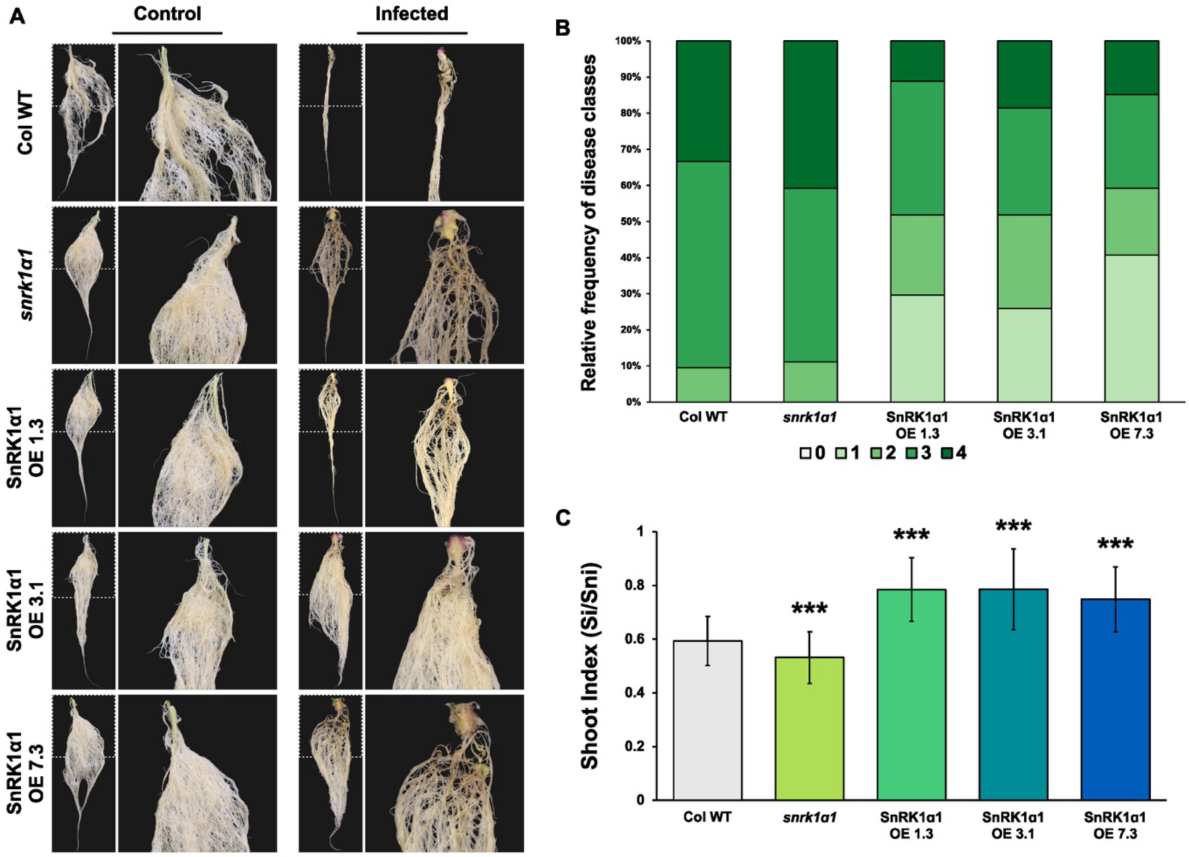


**Supplementary Figure 3. SnRK1α1 mediates resistance against clubroot disease development in Col-0 background plants in a miniature hydroponics setup.** (A) Representative root phenotypes of soil-grown infected and uninfected (control) wild-type *A. thaliana* (Col-0 WT), three independent SnRK1α1 overexpression lines (OE 1.3, OE 3.1, and OE 7.3), and a single *snrk1α1* T-DNA line grown in a miniature hydroponic system. (B)

Relative frequency distribution of clubroot disease severity classes among the representative SnRK1α1 lines. Symptom severity was scored based on five distinct classes: Class 0, no visible symptoms; Class 1, minor swellings on minor and/or secondary roots while maintaining typical root structure; Class 2, visibly thickened primary roots, reduced fine roots and lateral roots; Class 3, significantly reduced root system with visible galls on primary and secondary roots, loss of fine roots, and occasional gall development on the hypocotyl; Class 4, roots primarily composed of a single sizeable brownish gall. (C) Relative shoot weight (shoot index, SI) was calculated for each line as the ratio of the fresh weight of infected plants to that of non-infected control plants (Si/Sni). Fourteen-day-old *A. thaliana* seedlings were inoculated with *P. brassicae* 'e3' single spore suspension ($10^9$ spores/mL in 50 mM $KH_2PO_4$, pH 5.5), and disease progress was evaluated at 28 days post-inoculation (dpi). Data are the mean ± SD of three independent biological replicates, a total of 60–80 plants. Error bars represent the standard error of the means of the three replicates. Asterisks indicate a significant difference compared with the wild-type (for *** $p < 0.001$; two-way analysis of variance).

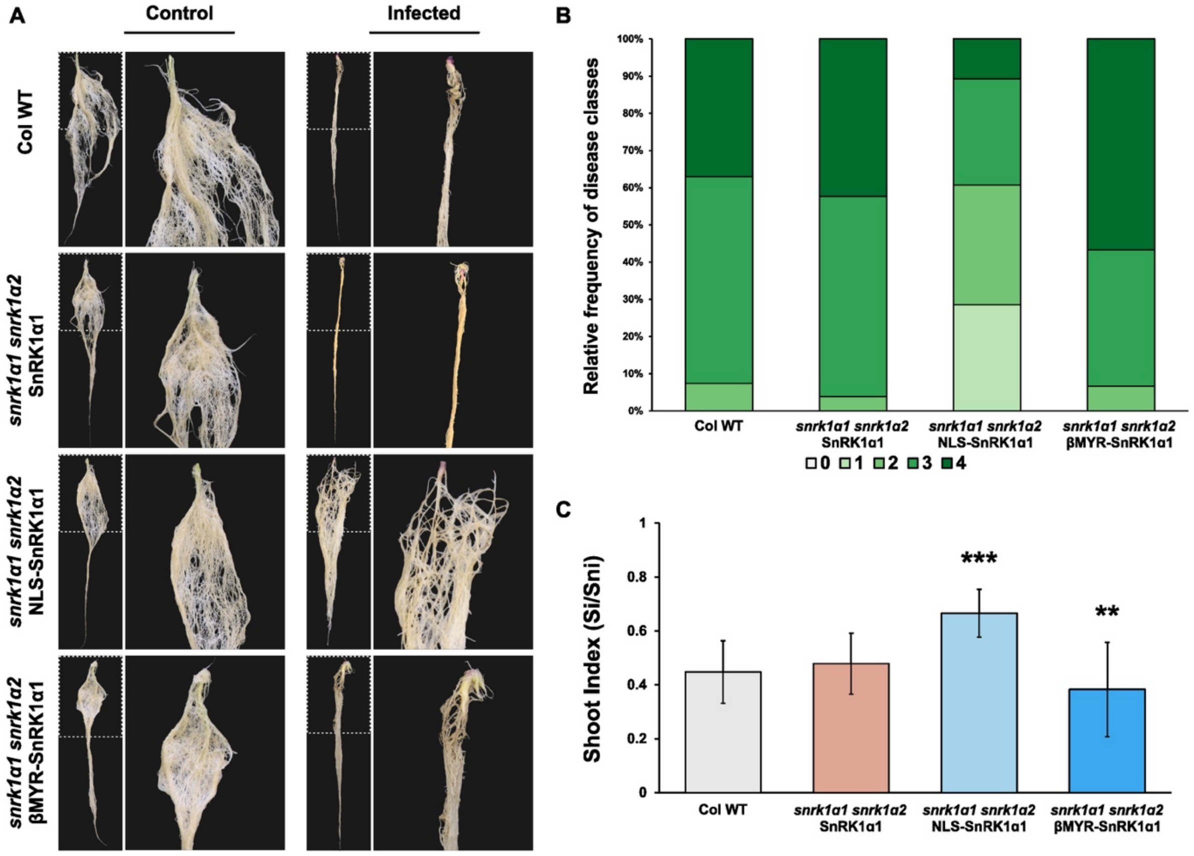


**Supplementary Figure 4. Increased nuclear SnRK1α1 localization increases resistance against clubroot disease in a miniature hydroponics setup.** (A) Representative root phenotypes of infected and uninfected (control) wild-type *A. thaliana* (Col-0 WT) and *snrk1α1 snrk1α2* double KO mutants complemented with either genomic SnRK1α1 fragments (SnRK1α1), nuclear-localized SnRK1α1 (NLS-SnRK1α1), or cytoplasmic retained SnRK1α1 (βMYR-SnRK1α1), grown in a miniature hydroponic system. (B) Relative frequency distribution of clubroot disease severity classes among the representative SnRK1α1 lines. Symptom severity was scored based on five distinct classes: Class 0, no visible symptoms; Class 1, minor swellings on minor and/or secondary roots while maintaining typical root structure; Class 2, visibly thickened primary roots, reduced fine roots and lateral roots; Class 3, significantly reduced root system with visible galls on primary and secondary roots, loss of fine roots, and occasional gall development on the hypocotyl; Class 4, roots primarily composed of a single sizeable brownish gall. (C) Relative shoot weight (shoot index, SI) was calculated for each line as the ratio of the fresh weight of infected plants to that of non-infected control plants (Si/Sni). Fourteen-day-old *A. thaliana* seedlings were inoculated with *P. brassicae* 'e3' single spore suspension ($10^9$ spores/mL in 50 mM $KH_2PO_4$, pH 5.5), and disease progress was evaluated at 28 days post-inoculation (dpi). Data are the mean ± SD of three independent biological replicates, a total of 60–80 plants. Error bars represent the standard error of the means of the three replicates. Asterisks indicate a significant difference compared with the wild-type (for ** $p < 0.01$; *** $p < 0.001$; two-way analysis of variance).

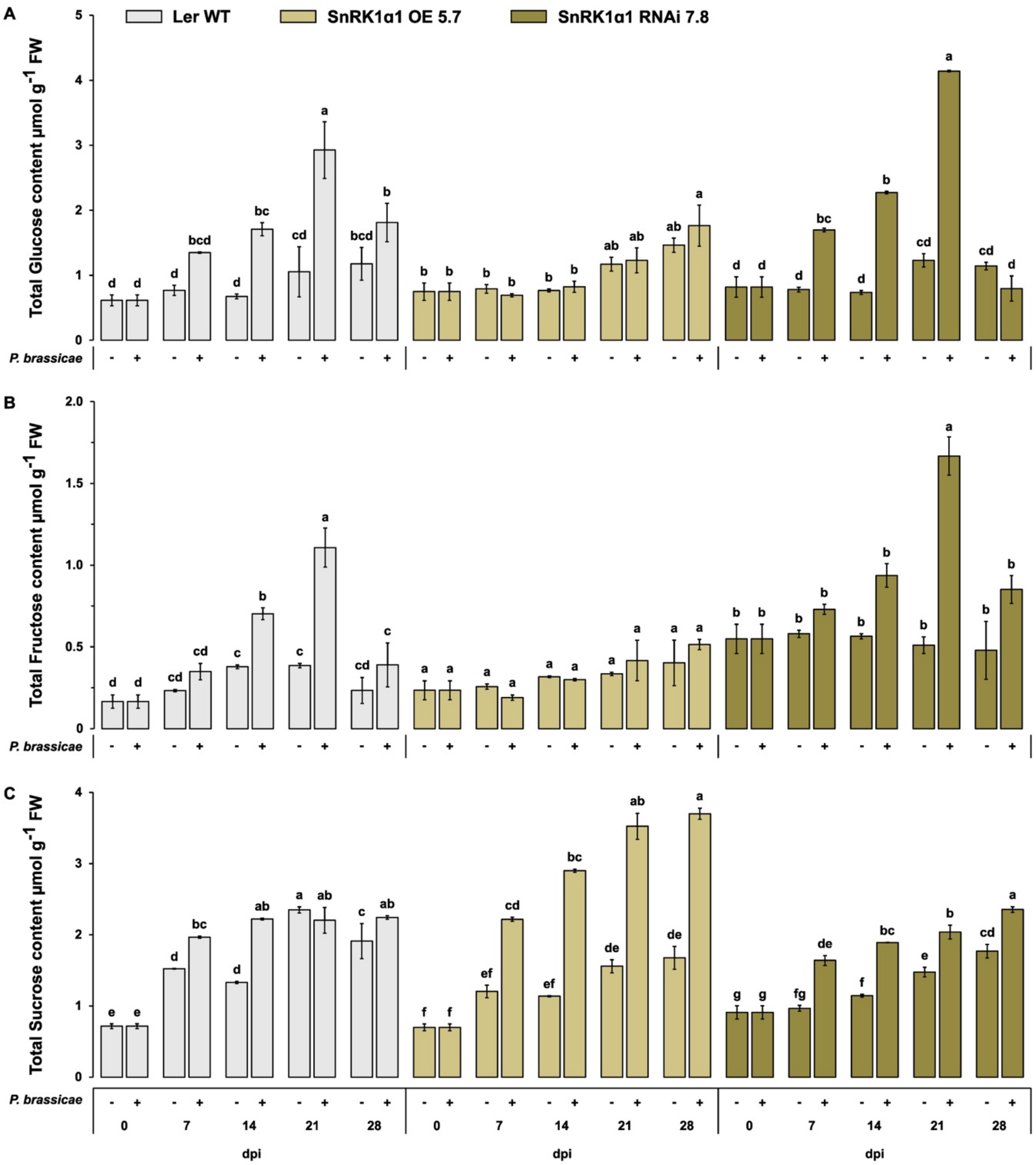


**Supplementary Figure 5. SnRK1α1 overexpression limits the impact of *P. brassicae* infection on soluble sugar dynamics in *A. thaliana* Ler-0 ecotype roots.** Soluble sugar content (A) Glucose, (B) Fructose, and (C) Sucrose was measured in non-infected and infected roots of different SnRK1α1 modified-lines from hydroponically grown *A. thaliana* at 0, 7, 14, 21, and 28 dpi using HPAEC-IPAD. Data are the mean ± SD of three independent biological replicates, a total of 60–80 plants. Error bars represent the standard error of the means of the three replicates. Different letters represent statistically significant differences within each genotype ($p < 0.005$; two-way analysis of variance).

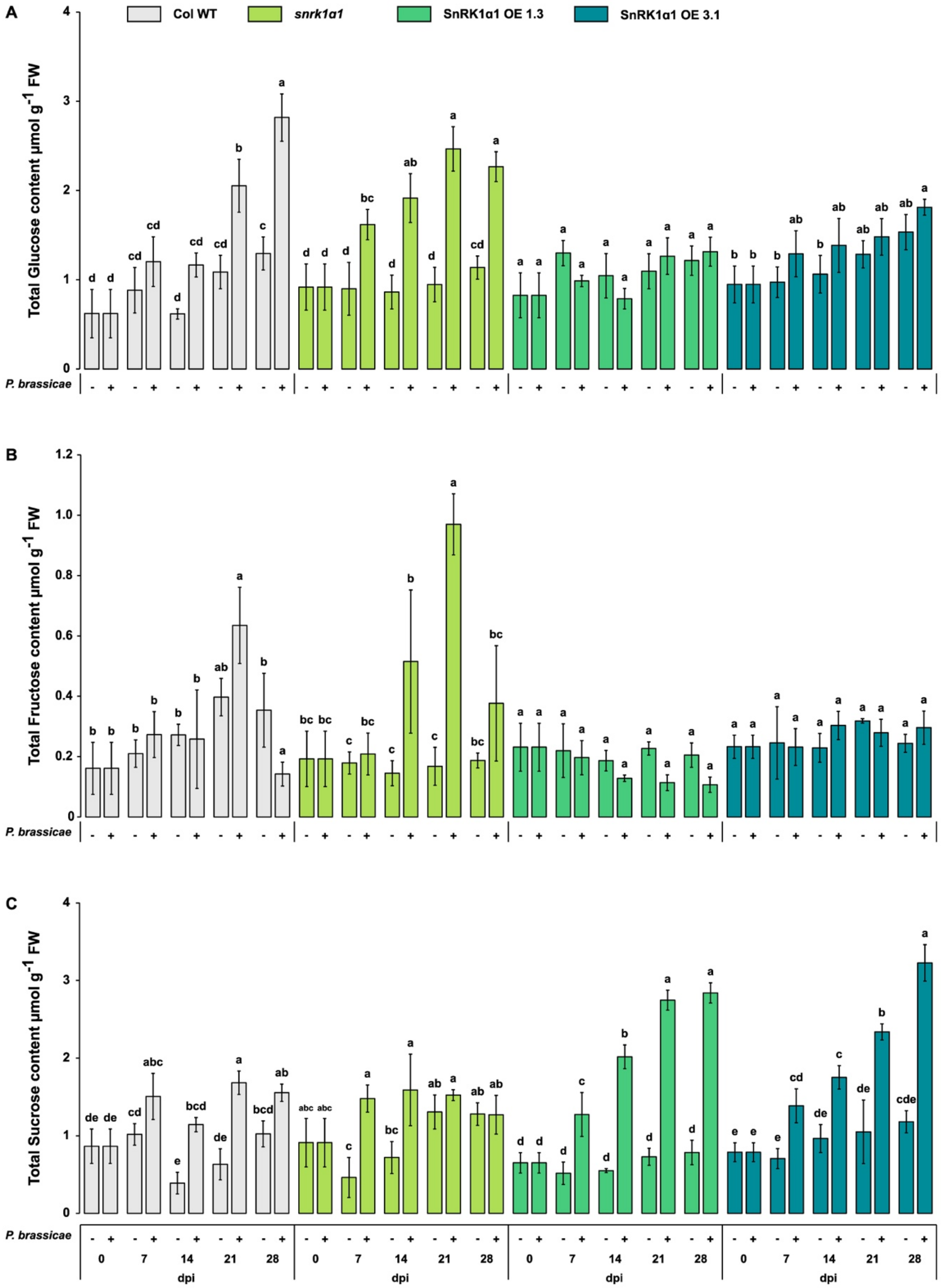


**Supplementary Figure 6. SnRK1α1 overexpression limits the impact of *P. brassicae* infection on soluble sugar dynamics in *A. thaliana* Col-0 ecotype roots.** Soluble sugar content (A) Glucose, (B) Fructose, and (C) Sucrose was measured in non-infected and infected roots of different SnRK1α1 modified-lines from hydroponically grown *A. thaliana* at 0, 7, 14, 21, and 28 dpi using HPAEC-IPAD. Data are the mean ± SD of three independent biological replicates, a total of 60–80 plants. Error bars represent the standard error of the means of the three replicates. Different letters represent statistically significant differences within each genotype ($p < 0.005$; two-way analysis of variance).

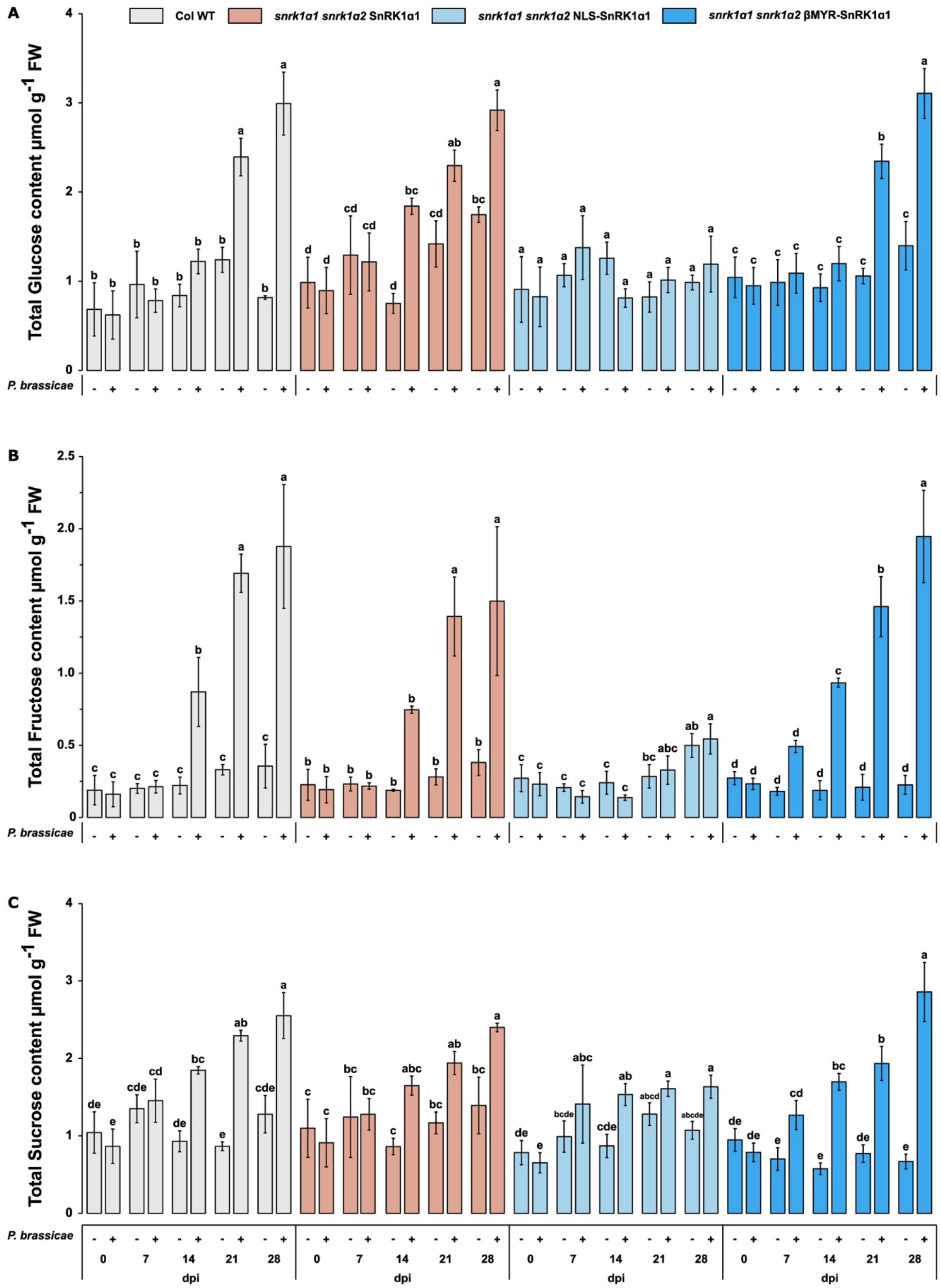


**Supplementary Figure 7. Increased nuclear SnRK1α1 localization limits the impact of *P. brassicae* infection on soluble sugar dynamics in *A. thaliana* roots.** Soluble sugar content (A) Glucose, (B) Fructose, and (C) Sucrose was measured in non-infected and infected roots of different SnRK1α1 modified-lines from hydroponically grown *A. thaliana* at 0, 7, 14, 21, and 28 dpi using HPAEC-IPAD. Data are the mean ± SD of three independent biological replicates, a total of 60–80 plants. Error bars represent the standard error of the means of the three replicates. Different letters represent statistically significant differences within each genotype ($p < 0.005$; two-way analysis of variance).

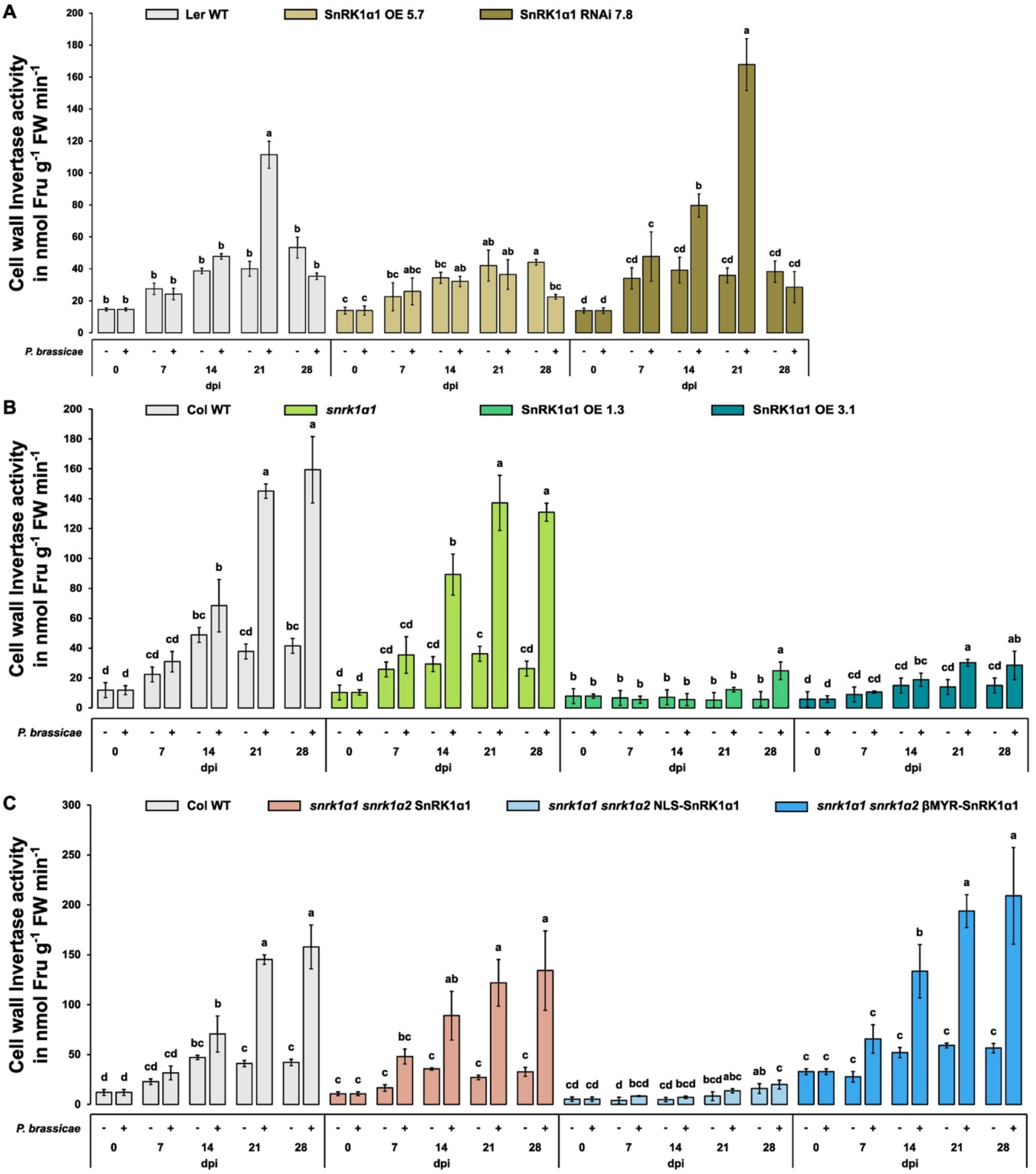


**Supplementary Figure 8. Increased nuclear SnRK1 reduces cell wall invertase (CWINV) activity in *A. thaliana* roots during clubroot infection.** Cell wall invertase activity was measured in non-infected and infected roots of (A-C) different SnRK1α1 modified-lines from hydroponically grown *A. thaliana* at 0, 7, 14, 21, and 28 dpi using HPAEC-IPAD. Enzyme activity was determined by quantifying fructose production from sucrose (nmol $min^{-1}$ $g^{-1}$ FW). Data are the mean ± SD of three independent biological replicates, a total of 60–80 plants. Error bars represent the standard error of the means of the three replicates. Different letters represent statistically significant differences within each genotype ($p < 0.005$; two-way analysis of variance).

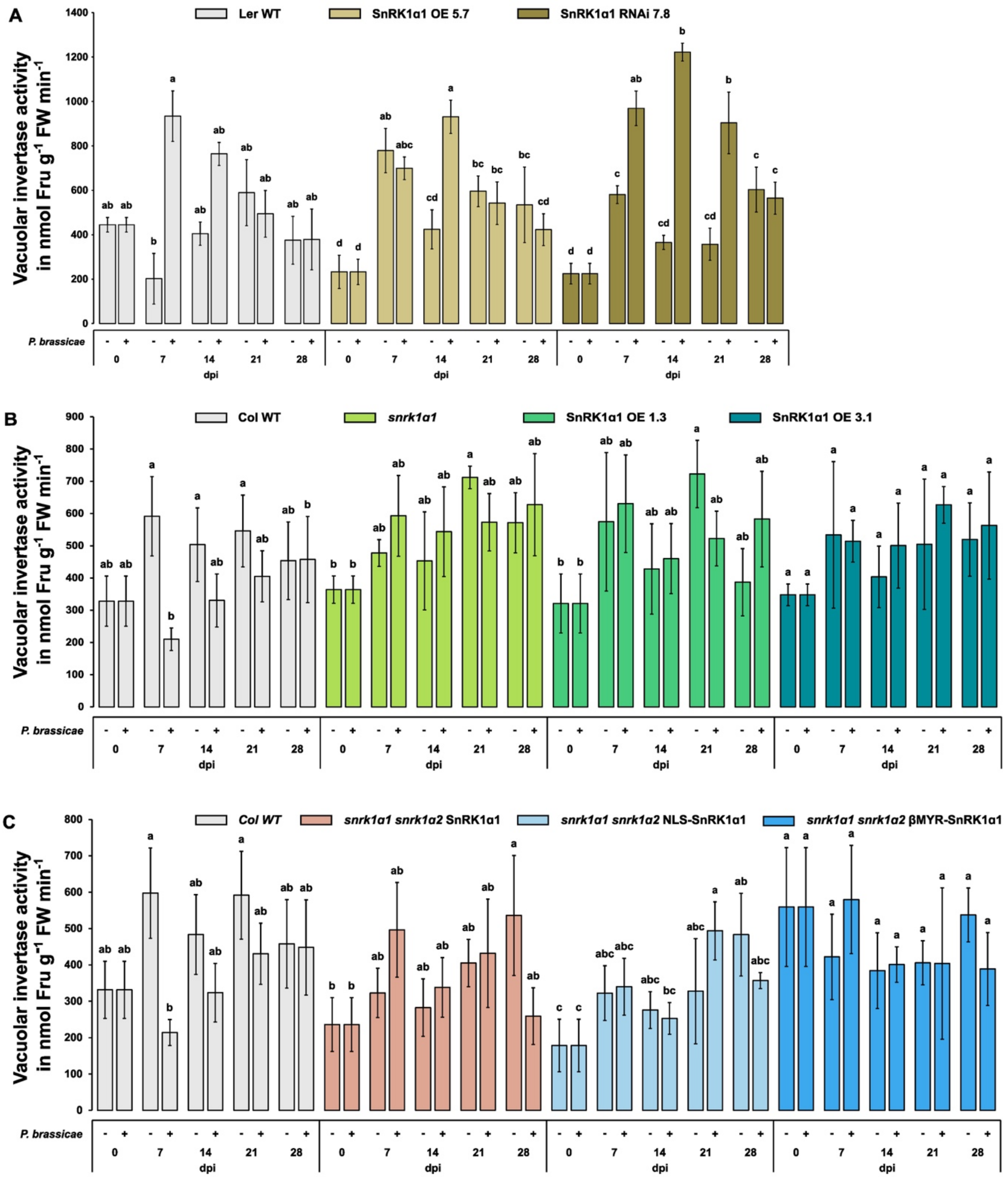


**Supplementary Figure 9. *P. brassicae* infection increases vacuolar invertase activity (VINV) in *A. thaliana* roots.** Vacuolar invertase activity was measured in non-infected and infected roots of (A-C) different SnRK1α1 modified-lines from hydroponically grown *A. thaliana* at 0, 7, 14, 21, and 28 dpi using HPAEC-IPAD. Enzyme activity was determined by quantifying fructose production from sucrose (nmol min$^{-1}$ g$^{-1}$ FW). Data are the mean ± SD of three independent biological replicates, a total of 60–80 plants. Error bars represent the standard error of the means of the three replicates. Different letters represent statistically significant differences within each genotype ($p < 0.005$; two-way analysis of variance).

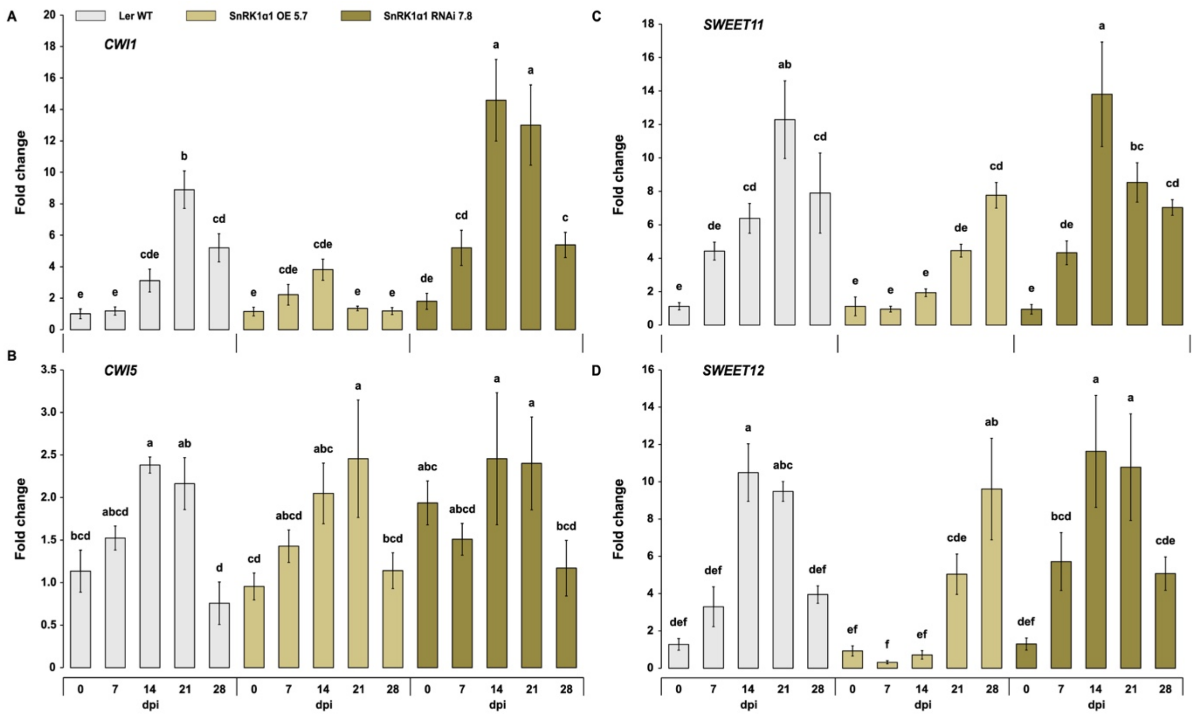


**Supplementary Figure 10. Increased nuclear SnRK1 activity limits the induction of cell wall invertases (*CWI1* & *CWI5*) and sugar transporters (*SWEET11* & *SWEET12*) in Ler-0 ecotype roots during clubroot infection.** (A-B) Cell wall invertase (*CWI1* & *CWI5*) expression and (C-D) expression of Sugars Will Eventually Be Exported Transporters (*SWEET11* & *SWEET12*) were measured in infected roots of different SnRK1α1 modified-lines from hydroponically grown *A. thaliana* at 0, 7, 14, 21, and 28 dpi using qRT-PCR. Data are the mean ± SD of three independent biological replicates, a total of 60–80 plants. Relative expression was normalized with the expression of *AtTUB* and calculated against 0 dpi. Error bars represent the standard error of the means of the three replicates. Different letters represent statistically significant differences across different genotypes ($p < 0.005$; two-way analysis of variance).

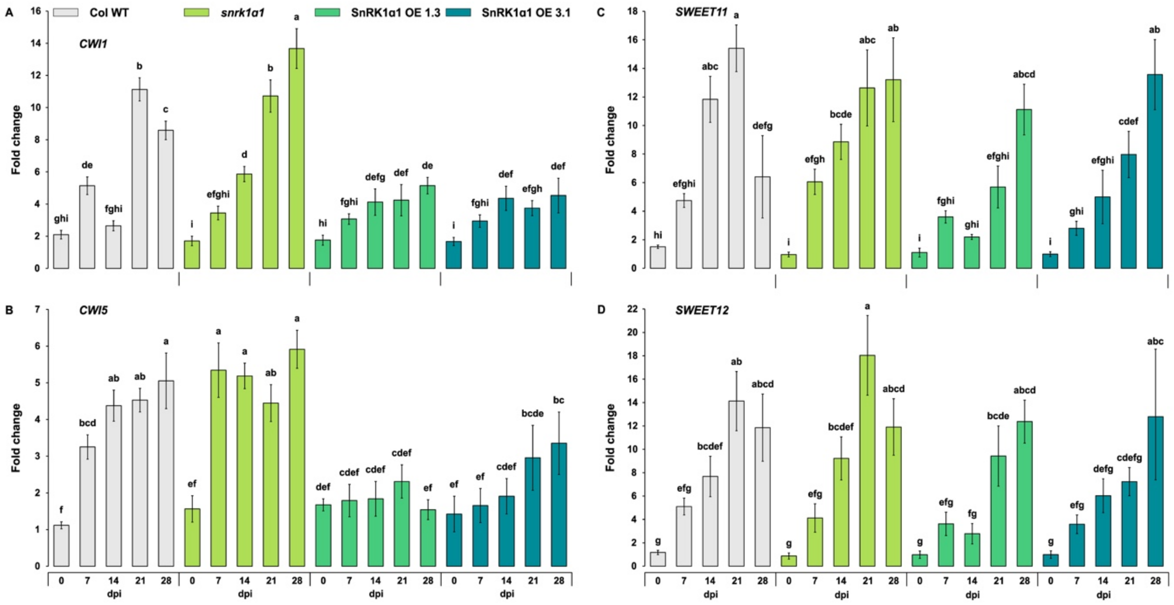


**Supplementary Figure 11. Increased SnRK1 activity limits the induction of cell wall invertases (*CWI1* & *CWI5*) and sugar transporters (*SWEET11* & *SWEET12*) in Col-0 ecotype roots during clubroot infection.** (A-B) Cell wall invertase (*CWI1* & *CWI5*) expression and (C-D) expression of Sugars Will Eventually Be Exported Transporters (*SWEET11* & *SWEET12*) were measured in infected roots of different SnRK1α1 modified-lines from hydroponically grown *A. thaliana* at 0, 7, 14, 21, and 28 dpi using qRT-PCR. Data are the mean ± SD of three independent biological replicates, a total of 60–80 plants. Relative expression was normalized with the expression of *AtTUB* and calculated against 0 dpi. Error bars represent the standard error of the means of the three replicates. Different letters represent statistically significant differences across different genotypes ($p < 0.005$; two-way analysis of variance).

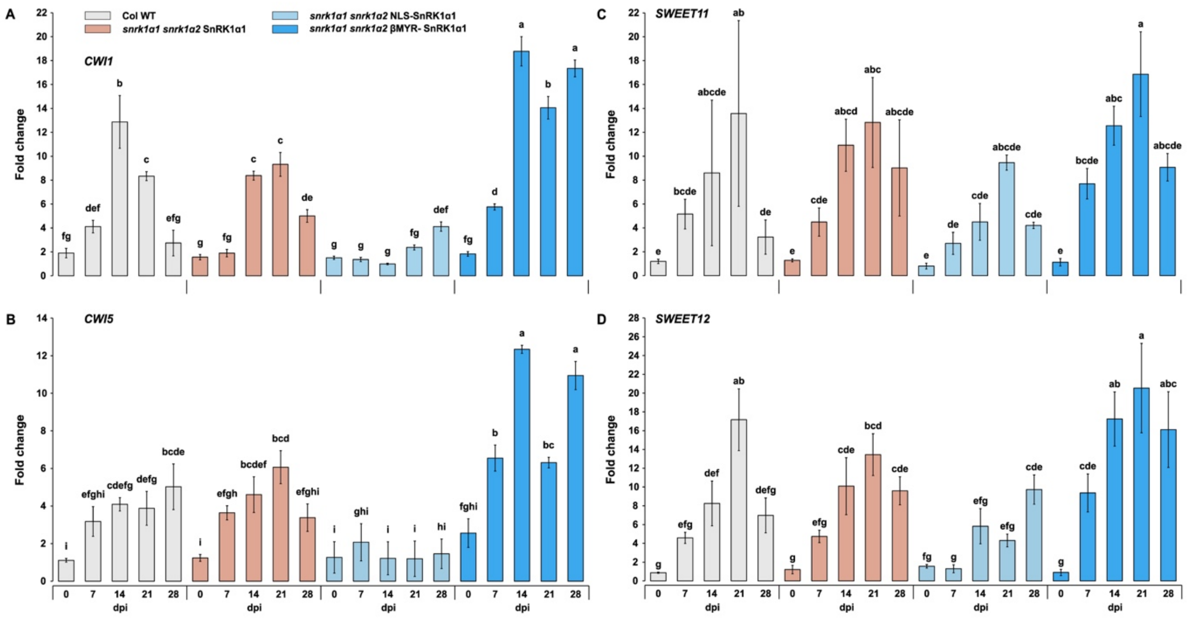


**Supplementary Figure 12. Increased nuclear SnRK1 activity limits the induction of cell wall invertases (*CWI1* & *CWI5*) and sugar transporters (*SWEET11* & *SWEET12*) in Col-0 ecotype roots during clubroot infection.** (A-B) Cell wall invertase (*CWI1* & *CWI5*) expression and (C-D) expression of Sugars Will Eventually Be Exported Transporters (*SWEET11* & *SWEET12*) were measured in infected roots of different SnRK1α1 modified-lines from hydroponically grown *A. thaliana* at 0, 7, 14, 21, and 28 dpi using qRT-PCR. Data are the mean ± SD of three independent biological replicates, a total of 60–80 plants. Relative expression was normalized with the expression of *AtTUB* and calculated against 0 dpi. Error bars represent the standard error of the means of the three replicates. Different letters represent statistically significant differences across different genotypes ($p < 0.005$; two-way analysis of variance).

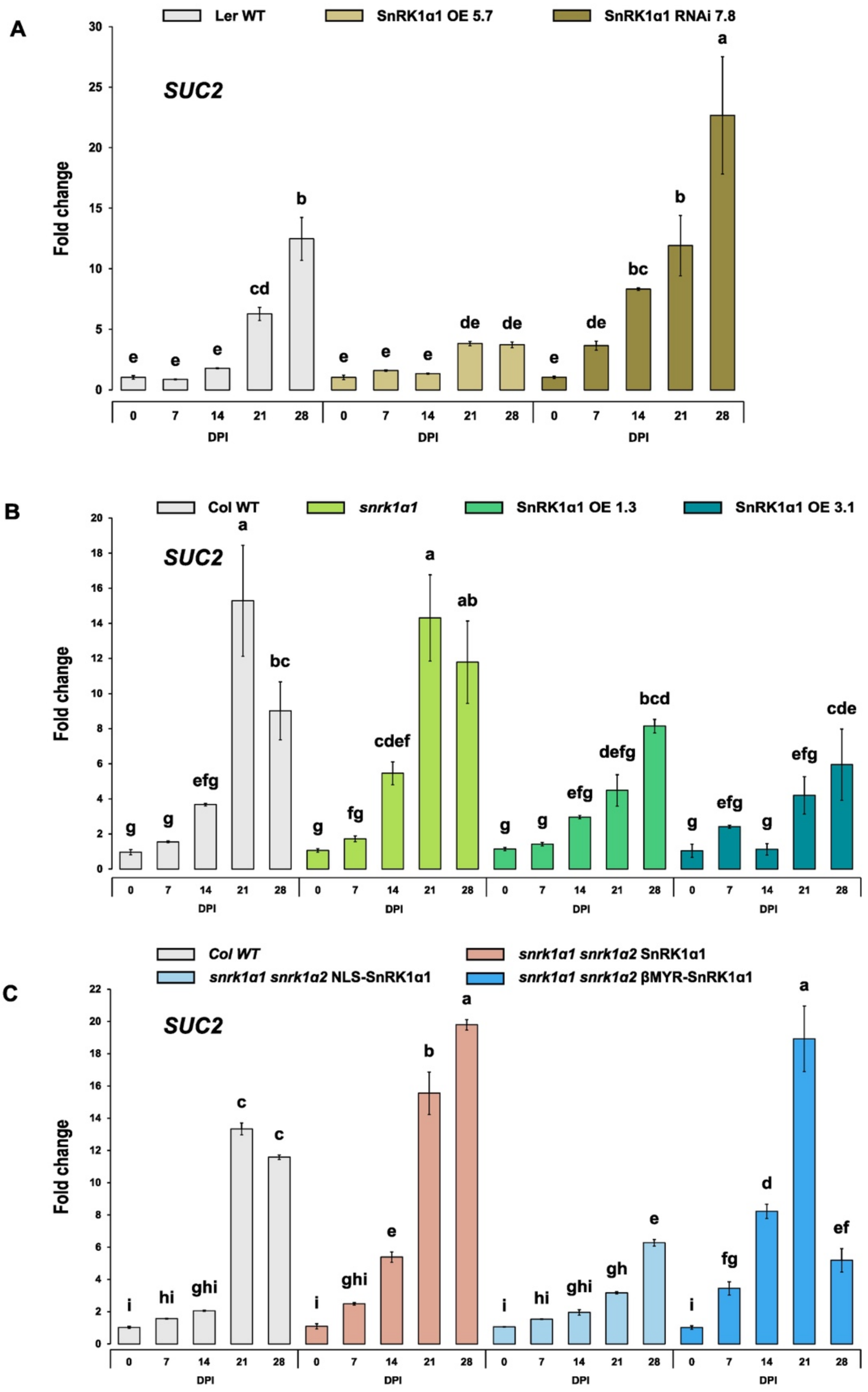


**Supplementary Figure 13. Increased nuclear SnRK1 activity limits the induction of Sucrose transporter 2 (SUC2) in clubroot-infected roots.** (A) SnRK1-modified plants in Ler-0 ecotype (B) SnRK1-modified plants in Col-0 ecotype (C) Altered SnRK1 localization lines in Col-0 ecotype. *SUC2* gene expression were measured in the infected roots from hydroponically grown *A. thaliana* at 0, 7, 14, 21, and 28 dpi using qRT-PCR. Data are the mean ± SD of three independent biological replicates, a total of 60–80 plants. Relative expression was normalized with the expression of *AtTUB* and calculated against 0 dpi. Error bars represent the standard error of the means of the three replicates. Different letters represent statistically significant differences across different genotypes ($p < 0.005$; two-way analysis of variance).

A

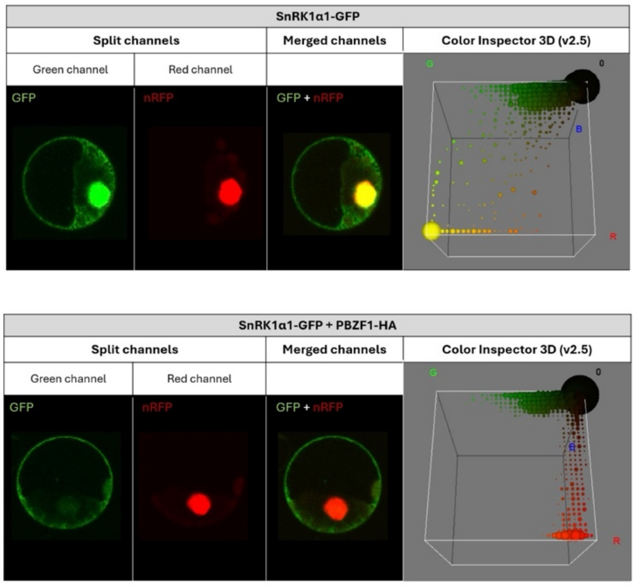


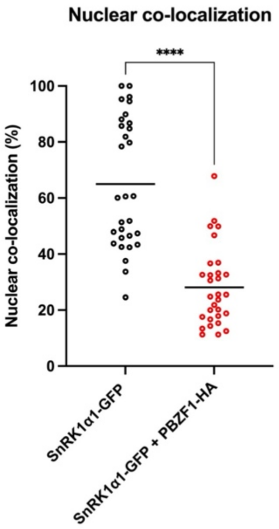


B

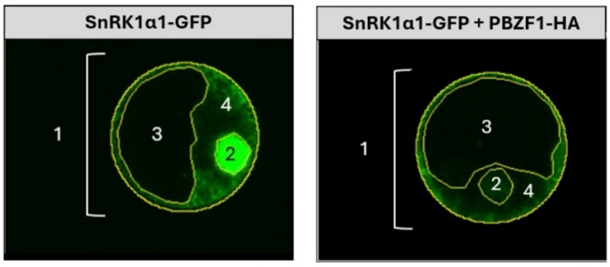


**SnRK1α1-GFP**

| | Area | Mean |
|---|---|---|
| 1 | 735.892 | 16.494 |
| 2 | 38.442 | 80.114 |
| 3 | 342.844 | 5.329 |
| 4 | 354.606 | 20.391 |

$$\frac{80.114}{20.391} = 3.93\ (> 1)$$

(increased **nuclear** SnRK1α1-GFP)

**SnRK1α1-GFP + PBZF1-HA**

| | Area | Mean |
|---|---|---|
| 1 | 796.712 | 8.958 |
| 2 | 38.768 | 12.821 |
| 3 | 463.941 | 4.608 |
| 4 | 294.003 | 15.313 |

$$\frac{12.821}{15.313} = 0.84\ (< 1)$$

(increased **cytoplasmic** SnRK1α1-GFP)

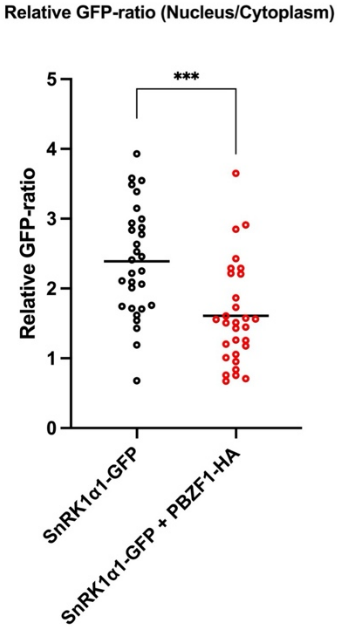


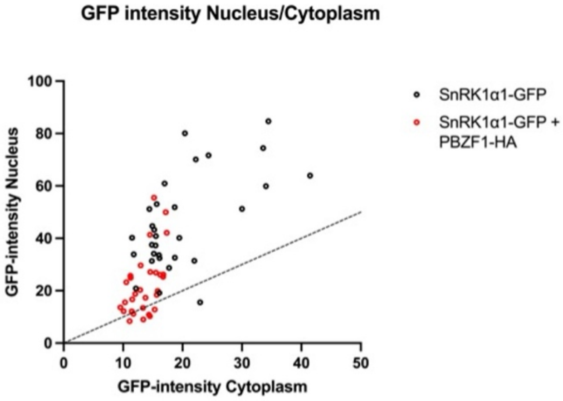

**Supplementary Figure 14. The PBZF1 effector affects the nuclear localization and relative nuclear-to-cytoplasmic GFP-ratio of SnRK1α1 in Arabidopsis (Col-0) mesophyll protoplasts.** (A) Nuclear marker co-localization and (B) relative nuclear-to-cytoplasmic GFP-ratio of SnRK1α1-GFP was measured in the presence or absence of PBZF1 14h after transfection using confocal fluorescence microscopy (FluoViewTM FV1000 confocal system). An SCF30-RFP (nRFP) construct was co-expressed and served as a red fluorescent nuclear marker. Confocal images from 30 individual protoplasts per condition were analyzed using Fiji (ImageJ). Values are averages with standard deviations, n=30. One-way ANOVA statistical analysis was performed, *** $P<0.001$, **** $P<0.0001$.

**Supplementary Table S1. Cloning and qRT-PCR primers.**

**Cloning Primers**

| **Plasmodiophora brassicae genes** | | | |
|---|---|---|---|
| Gene name | Gene ID | Forward primer | Reverse primer |
| PBZF1 | PBCN_001987 | CGGGATCCATGGCGCGATTCGCATGGTC | AAGGCCTGCTGTTGAAGCTCGTGCCAG |
| SnRK1α1 | AT3G01090 | CG-GGATCC-ATGGATGGATCAGGCACAGG | A-AGGCCT-GAGGACTCGGAGCTGAGCA |
| NLS-SnRK1α1 | AT3G01090 | CG-GGATCC-ATG-CCTCCAAAAAAGAAGAGAAAGGTAGCT-GATGGATCAGGCACAGGCAG | A-AGGCCT-GAGGACTCGGAGCTGAGCA |

**qRT-PCR Primers**

| **Arabidopsis genes** | | | |
|---|---|---|---|
| Gene name | Gene ID | Forward primer | Reverse primer |
| AtCWI1 | AT3G13790 | TTGAAGCCTCGCACCATGT | CATAGGCCCATTAGGATCGTT |
| AtCWI5 | AT3G13784 | TTGGATGAACGATCCTAACGGG | CGAGATGGTCTGATTGCTGGA |
| AtSWEET11 | AT3G48740 | TCCTTCTCCTAACAACTTATATACCATG | TCCTATAGAACGTTGGCACAGGA |
| AtSWEET12 | AT5G23660 | AAAGCTGATATCTTTCTTACTACTTCGAA | CTTACAAATCCTATAGAACGTTGGCAC |
| AtSUC2 | AT1G22710 | ACGAACTATTCGGTGGTGGAA | CTGCCGCAATCGCTCCTAA |
| AtTUB | AT5G44340 | AGGGAAACGAAGACAGCAAG | GCTCGCTAATCCTACCTTTGG |

| **Plasmodiophora brassicae genes** | | | |
|---|---|---|---|
| Gene name | Gene ID | Forward primer | Reverse primer |
| PbITS | Y12831 | TACCATACCCAGGGCGATT | CAACGAGTCAGCTTGAATGC |
| PbActin | AY452179 | CGGGACATCACCGACTACCT | TGGACATCTCGGTGTCGAAGT |
